# Loss of Neurofibromin Induces Inflammatory Macrophage Phenotypic Switch and Retinal Neovascularization via GLUT1 Activation

**DOI:** 10.1101/2024.09.13.612509

**Authors:** Yusra Zaidi, Rebekah Tritz, Nida Zaidi, Faisal Nabi, Syed Adeel H. Zaidi, Abdelhakim Morsy, Valerie Harris, Rilee Racine, Farlyn Z. Hudson, Zsuzsanna Bordan, Simone Kennard, Robert Batori, Yuqing Huo, Gabor Csanyi, Eric J. Belin de Chantemèle, Kecheng Lei, Nicholas M. Boulis, David J. Fulton, Rizwan Hasan Khan, Ruth B. Caldwell, Brian K. Stansfield

## Abstract

Persons with neurofibromatosis type 1 (NF1), a tumor predisposition syndrome, are largely protected from diabetes and exhibit evidence of enhanced glucose metabolism, which is replicated in mice harboring *Nf1* mutations. A hallmark of NF1-associated neurofibromas and sarcomas is the high density of inflammatory macrophages and targeting macrophages appears efficacious in models of NF1. Inflammatory macrophages rely on glycolysis to rapidly generate ATP; thus, identifying whether neurofibromin, the protein encoded by the *NF1* gene, controls glucose uptake and/or glycolysis in macrophages is therapeutically compelling. Using neurofibromin-deficient macrophages and macrophage-specific *Nf1* knockout mice, we demonstrate that neurofibromin complexes with glucose transporter 1 (GLUT1) to restrain its activity and that loss of neurofibromin permits Akt2 to facilitate GLUT1 translocation to the membrane in macrophages. In turn, glucose internalization and glycolysis are highly up regulated and provoke putative reparative (M2) macrophages to undergo inflammatory phenotypic switch. Inflammatory M1 macrophages and inflammatory-like M2 macrophages invest the perivascular stroma of tumors and induce pathologic angiogenesis in mice harboring macrophage-specific *Nf1* deletion. These studies identify a clear mechanism for the enhanced glycolysis and low risk for diabetes observed in persons with NF1 and provide a novel therapeutic target for manifestations of NF1.

## Introduction

Neurofibromatosis type 1 (NF1) is a common tumor predisposition syndrome affecting 1 in 2,500 persons worldwide(1). Mutations in the *NF1* gene, which causes NF1, have been identified in each of the 60 exons with more than 2,800 mutations identified to date(2). The high mutation rate and size of the *NF1* gene disfavor strong genotype-phenotype correlations. One clear exception to this rule is the highly conserved anthropometric and metabolic phenotype featuring short stature, low body mass index (BMI), and protection from diabetes, which has been described in multiple NF1 cohorts and registry studies(3–6). For example, a recently published registry study comparing 1,400 persons with NF1 to 14,000 matched controls revealed a 75% reduction in diabetes prevalence in persons with NF1(7). Strikingly, the NF1 cohort was similarly protected from diabetes when compared with non-NF1 siblings suggesting that *NF1* mutations, but not other genetic or environmental factors, convey this metabolic phenotype. Recently, we demonstrated that mice harboring germline mutations in a single *Nf1* allele (*Nf1*+/−) recapitulate many of the metabolic features observed in persons with NF1(8). *Nf1*+/− mice rapidly metabolize glucose and are protected from diet-induced hyperglycemia and obesity when compared with littermate WT mice; however, the response to insulin and glucagon did not differ between genotypes suggesting that neurofibromin, the protein product of *NF1*, controls basal glucose uptake and/or glycolysis. The mechanism through which neurofibromin controls glucose metabolism is unknown.

The increase in basal glucose utilization suggested from observational studies in persons with NF1 and in experimental studies in *Nf1*-mutant mice might be leveraged for novel therapeutics. In this regard, macrophages are particularly attractive targets since functional macrophage populations rely on different metabolic substrates with “inflammatory” macrophages tending to use glycolysis for rapid generation of ATP(9). This rationale is in keeping with the emergence of clinical and experimental data demonstrating that persons with NF1 and mice harboring *Nf1* mutations experience chronic inflammation characterized by increased frequency of circulating inflammatory monocytes and cytokines as well as enhanced free radical formation(10–15). Macrophages comprise nearly 40% of the total cell mass of NF1 tumors and targeting macrophage-mediated byproducts has demonstrated efficacy in reducing tumor size, slowing malignant transformation, suppressing arteriopathy, and promoting healthy bone resorption in experimental models of NF1. In many of these models, macrophage subtyping reveals that inflammatory macrophages are the most common population, although other macrophage subpopulations are present, and accumulate in the peri-vascular stroma of tumors and in the media of arteries(15–17). These observations may be useful for understanding the role of neurofibromin in controlling basal glucose metabolism and macrophage differentiation based on recent data describing enhanced carbohydrate metabolism in persons with NF1 and in animals harboring *Nf1* mutations. Whether this highly conserved phenotype may be leveraged for understanding the role of neurofibromin in macrophage polarization and function has not been explored.

Here, we provide genetic and biochemical evidence that neurofibromin interacts with glucose transporter 1 (GLUT1) to maintain GLUT1 translocation to the plasma membrane of macrophages. Loss of neurofibromin, as seen in persons with NF1, increases membrane-bound GLUT1 mediated glucose uptake to promote putative reparative (M2) macrophages to undergo phenotypic switch to an inflammatory phenotype, which promotes pathologic retinal neovascularization.

## Results

### Loss of neurofibromin enhances glucose utilization in macrophages

Macrophages undergo metabolic reprogramming as they differentiate into inflammatory and reparative phenotypes, or so-called M1 and M2 populations, respectively. We derived macrophages from the long bones of *Nf1^flox/flox^*;LysMCre^+^ (*Nf1*^ΔMΦ^) and *Nf1^flox/flox^*;LysMCre^-^ (*Nf1*^fl/fl^) mice and exposed macrophages to lipopolysaccharide (LPS) and interferon gamma (IFNγ) to induce an inflammatory (M1) phenotype or interleukin-4 (IL4) to induce a reparative (M2) phenotype prior to metabolic characterization using the Seahorse XF Extracellular Flux Analyzer. First, we compared the metabolic potential between each *Nf1*^ΔMΦ^ and *Nf1*^fl/fl^ macrophage population before and after the simultaneous injection of oligomycin and Trifluoromethoxy carbonylcyanide phenylhydrazone (FCCP) to increase glycolysis and oxygen consumption simultaneously. While *Nf1*^ΔMΦ^ and *Nf1*^fl/fl^ macrophages exhibited the same basal metabolic phenotype, the metabolic potential of *Nf1*^ΔMΦ^ macrophages was exaggerated in some subpopulations (**Figure 1A – F**). Oxygen consumption rate (OCR) and extracellular acidification rate (ECAR), indicative of mitochondrial respiration and glycolysis respectively, were significantly higher in non-polarized (M0) and inflammatory (M1) *Nf1*^ΔMΦ^ macrophages, while no differences in metabolic potential were observed between genotypes of reparative (M2) macrophages. As seen in **Figure 1 (A-C)** the primary metabolic pathways for non-polarized and reparative MΦs are similar while inflammatory MΦs are highly reliant on glycolysis to meet their energy needs.

**Figure 1.**
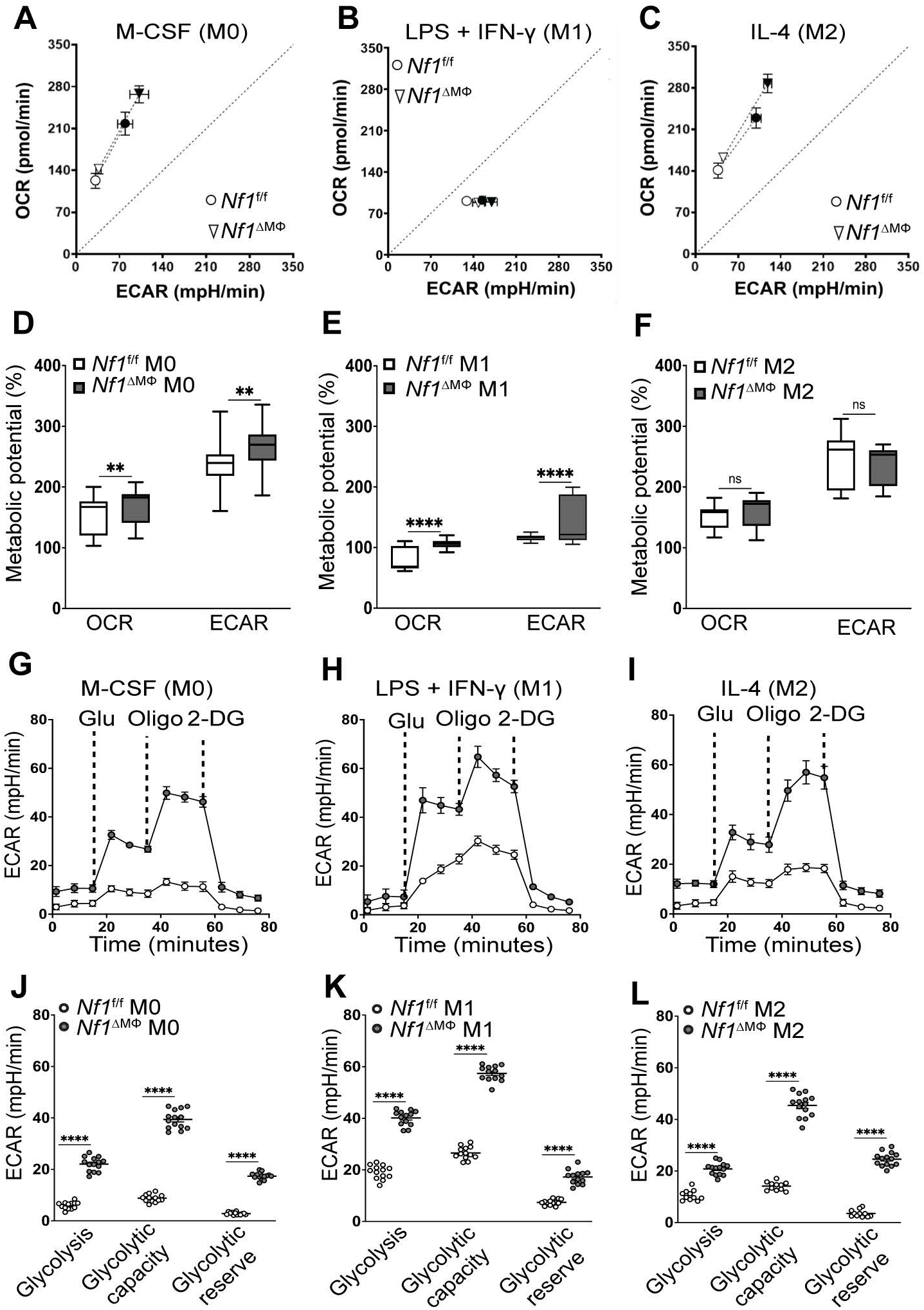
Loss of neurofibromin enhances glycolysis in macrophages. Representative measurements of **(A-F)** percent metabolic potential, and **(G-L)** Extracellular acidification rate (EGAR) performed on BMDMs derived from long bones of *Nf1*^f/f^ and *Nf1*^ΔMφ^ mice treated with 20ng/ml LPS and IFN-γ or 20ng/ml IL-4 for polarizing the macrophages to inflammatory (M1) or reparative (M2) for 16 hours, respectively. Subsequent addition of glucose, the ATP synthase inhibitor oligomycin, and the hexokinase inhibitor 2-deoxy-glucose (2-DG) were carried out where indicated **(G-I),** n = 4 mice/genotype, each data point represents technical replicate Data are expressed as mean ± SEM. P values were calculated using One-way ANOVA followed by Tukey’s multiple comparision test (*P < 0.05, ** **P** < 0.01,*** P < 0.001 **** P < 0.0001, ns, not significant).

Next, we performed a glycolysis stress test in *Nf1*^ΔMΦ^ and *Nf1*^fl/fl^ macrophages. In each macrophage population, *Nf1*^ΔMΦ^ macrophages displayed significantly higher rates of basal glycolysis, glycolytic capacity, and glycolytic reserve when compared with *Nf1*^fl/fl^ macrophages (**Figure 1G – L**). A striking feature of these studies was the high glycolytic capacity and reserve of reparative (M2) *Nf1*^ΔMΦ^ macrophages that approximated the values observed in the traditionally glycolytic inflammatory (M1) population of *Nf1*^ΔMΦ^ macrophages. We observed similar relationships in neurofibromin-deficient human THP1-derived macrophages (**Figures S1 & S2**). These observations were unique to neurofibromin-deficient MΦs and suggests that loss of neurofibromin enhances glycolysis independent of the polarizing agent.

After observing enhanced glycolysis in each *Nf1*^ΔMΦ^ macrophage population, we sought to identify whether the genotypic differences reflect higher substrate availability. Each *Nf1*^ΔMΦ^ and *Nf1*^fl/fl^ macrophage population was incubated with a glucose analog, 2-deoxy-2-[(7-nitro-2,1,3-benzoxadiazol-4-yl) amino]-D-glucose (2-NBDG) that is readily taken up by cells but fails to undergo the initial steps of glycolysis and accumulates in the cell. In the *Nf1*^fl/fl^ macrophage (i.e., control) populations, glucose uptake mirrored the trends observed in the glycolysis stress test wherein glucose uptake is modestly increased in inflammatory (M1) macrophages and similar glucose internalization was observed between non-polarized and reparative (M2) macrophages (**Figure 2A**). As seen in the glycolysis stress test, glucose uptake was significantly increased in all *Nf1*^ΔMΦ^ macrophage populations when compared with each *Nf1*^fl/fl^ macrophage population (**Figure 2A – B**). Most striking was the observation that glucose uptake in reparative (M2) *Nf1*^ΔMΦ^ macrophages approximated and exceeded the glucose uptake observed in inflammatory (M1) *Nf1*^ΔMΦ^ macrophages (**Figure 2B**). Glucose internalization was inhibited in all *Nf1*^ΔMΦ^ macrophage populations by BAY876, a specific inhibitor of glucose transporter 1 (GLUT1), suggesting that basal glucose uptake via GLUT1 accounts for the increase in intracellular glucose and resultant glycolysis observed in *Nf1*^ΔMΦ^ macrophages (**Figure 2C-E**).

**Figure 2.**
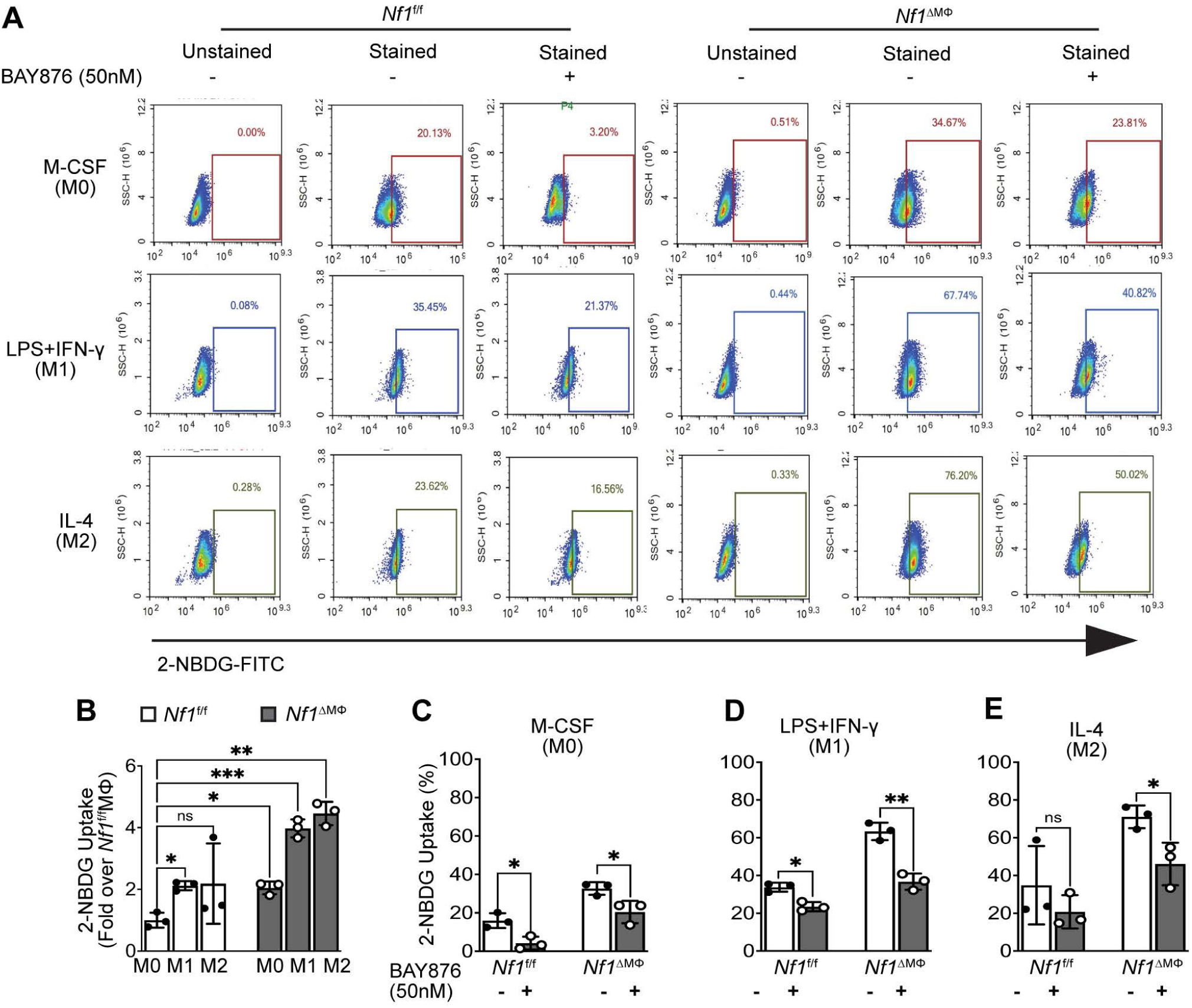
Glucose utilization is enhanced in neurofibromin-deficient macrophages via GLUT1. **(A)** Representative flow cytometric plots, and **(B-E)** quantification of percent glucose uptake by viable BMDMs derived from bones of *Nf1*^f/f^ and *Nf1*^ΔMφ^ mice in absence or presence of GLUT1 inihibitor, BAY876. BMDMs were labeled with 2-NBDG at a concentration of 100µM for 30 minutes in absence or presence of BAY876 (50nM), to measure glucose uptake by flow cytometry, n = 3 mice per genotype. Data are expressed as mean ± SD. P values were calculated using One-way ANOVA followed by Dunnette’s T3 multiple comparison test for **(B)** and 2-tailed Student’s t-test for (**C, D,** and **E**). (*P < 0.05, ** **P** < 0.01,*** P < 0.001 **** P < 0.0001, ns, not significant).

### Neurofibromin regulates GLUT1 localization to the plasma membrane

Glucose transporters (GLUT) are highly conserved, membrane-bound proteins with varied tissue expression, substrate affinity, and regulatory mechanisms. Within the GLUT family, GLUT1 is ubiquitously expressed, regulates basal glucose uptake independent of insulin and drives the pro-inflammatory phenotype of macrophages, ensuring sufficient glucose uptake (18–21). Based on our previous observations that enhanced glucose utilization in *Nf1*-mutant animals is insulin-independent (8) and present observations that glucose uptake and glycolysis are up-regulated in *Nf1*^ΔMΦ^ MΦs, we suspected that GLUT1 expression or function may be altered in *Nf1*^ΔMΦ^ MΦs. As seen in **Figure 3A**, GLUT1 is predominantly expressed in inflammatory *Nf1*^fl/fl^ macrophages and a similar, but exaggerated increase in GLUT1 was observed in inflammatory (M1) *Nf1*^ΔMΦ^ macrophages. Consistent with the increase in glucose uptake and glycolysis observed in reparative (M2) *Nf1*^ΔMΦ^ macrophages, we identified a significant increase in GLUT1 expression in reparative (M2) *Nf1*^ΔMΦ^ macrophages that was not indicative of *Nf1*^fl/fl^ reparative (M2) macrophages (**Figure 3A – C)** with confirmation at the mRNA level (**Figure 3D**). Next, we isolated the plasma membrane and cytosolic fractions from *Nf1*^ΔMΦ^ and *Nf1*^fl/fl^ macrophages and probed for GLUT1 expression. We observed an increase in GLUT1 in the plasma membrane of inflammatory (M1) *Nf1*^ΔMΦ^ macrophages when compared with inflammatory (M1) *Nf1*^fl/fl^ macrophages (**Figure 3E**). Consistent with our observations in the whole cell lysates, membrane bound GLUT1 was increased over 3-fold in reparative (M2) *Nf1*^ΔMΦ^ macrophages when compared with reparative (M2) *Nf1*^fl/fl^ macrophages (**Figure 3E – F**). Further, we observed a reciprocal decrease in cytosolic GLUT1 expression in reparative (M2) *Nf1*^ΔMΦ^ macrophages that was not observed in reparative (M2) *Nf1*^fl/fl^ macrophages (**Figure 3E & G**). Taken together, these data suggest that loss of neurofibromin facilitates GLUT1 translocation to the plasma membrane in reparative (M2) macrophages to mirror the glucose utilization phenotype traditionally observed in inflammatory (M1) macrophages.

**Figure 3.**
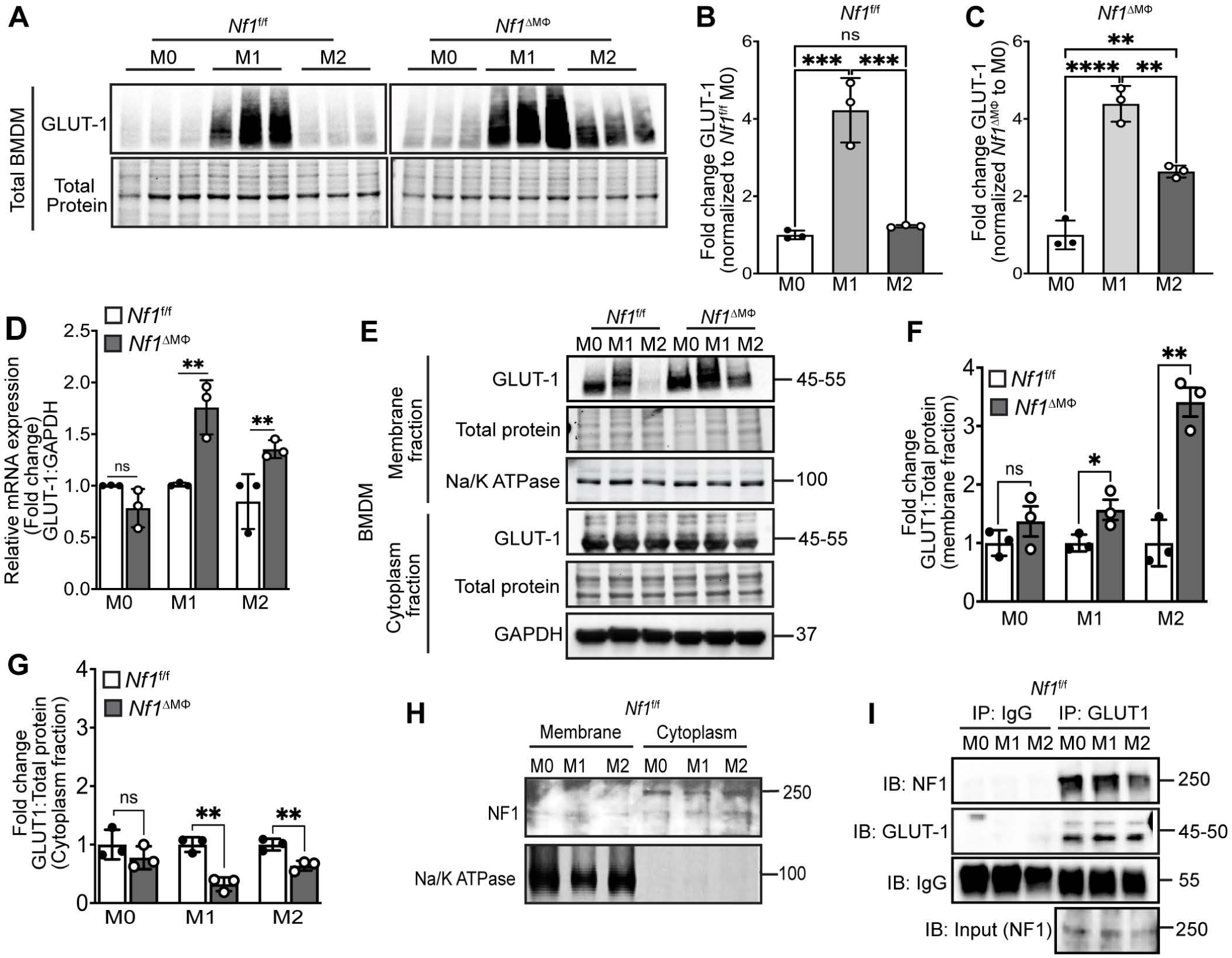
Loss of Neurofibromin increases facilitative GLUT1 expression in all macrophages. Representative immunoblots and quantification of GLUT1 expression in total BMDM lysates **(A, B** and **C)** and in membrane and cytoplasm fractions of BMDM **(E-G)** derived from bones of *Nf1*^f/f^ and *Nf1*^ΔMφ^ mice. BMDMs were differentiated using M-CSFand polarized to M1 and M2 macrophages, GLUT1 expression was measured by immunoblotting in all sub-popluation of macrophages, n=3 mice per genotype. **(D)** qRT-PCR analysis of GLUT1 mRNA relative expression in all subpopulation of macrophages (MO, M1, and M2) from bones of *Nf1*^f/f^ and *Nf1*^ΔMφ^ mice, n=3 mice per genotype, each data point represent the mean of two technical replicates of each mice. **(H)** Representative immunoblot of neurofibromin expression in membrane and cytoplasm frations of all macrophage sub-populations of *Nf1*^f/f^ mice. (I) Polarized BMDMs lysates from *Nf1*^f/f^ mice were prepared and GLUT1 was immunoprecipitated with anti-GLUT1 antibody. The immune complex was then immunoblotted with anti-NF1 antibody to identify GLTU1:NF1 complex. Data are expressed as mean ± SD. P values calculated using One-way ANOVA followed by Tukey’s multiple comparison test for **(B, C** and **D)** and 2-tailed Student’s t-test for **(F** and **G)** (*P < 0.05, ** P < 0.01, *** P < 0.001, **** P < 0.0001, ns, not significant).

Based on the increase in GLUT1 expression in the plasma membrane of *Nf1*^ΔMΦ^ macrophages, we fractionated *Nf1*^fl/fl^ macrophages to identify potential relationships between wildtype neurofibromin and GLUT1. We identified that neurofibromin is primarily expressed in the cytoplasm of all macrophage populations (**Figure 3H**). Based on this data and the increased expression of GLUT1 in *Nf1*^ΔMΦ^ macrophages, we hypothesized that neurofibromin interacts with GLUT1 in the cytoplasm of macrophages to restrain GLUT1 from localizing to the plasma membrane in *Nf1*^fl/fl^ macrophages. Therefore, the absence of neurofibromin would allow GLUT1 to localize to the plasma membrane. To interrogate this hypothesis, we precipitated GLUT1 from each *Nf1*^fl/fl^ macrophage population and probed for neurofibromin. As seen in **Figure 3I**, neurofibromin co-precipitates with GLUT1 in all *Nf1*^fl/fl^ macrophage populations. We conclude that loss of neurofibromin leads to unrestrained GLUT1 translocation to the plasma membrane of macrophages and permits glucose internalization and utilization.

### Loss of neurofibromin increases GLUT1 expression via Akt2

The increase in membrane bound GLUT1 in the absence of neurofibromin explains the enhanced glucose uptake and glycolysis observed in *Nf1*^ΔMΦ^ macrophages, but we also observed an increase in the expression of GLUT1 in whole *Nf1*^ΔMΦ^ macrophage lysates. Neurofibromin is principally recognized as a regulator of the Ras-Mek-Erk pathway, which is the focus of several inhibitor agents for persons with NF1, but Ras also regulates pathways linked to GLUT expression such as PI3-kinase and Akt. Therefore, we probed whole cell lysates from each *Nf1*^ΔMΦ^ and *Nf1*^fl/fl^ macrophage population and identified an activation of PI3-kinase (Phosphorylated PI3K) and both Akt isoforms in inflammatory (M1) and reparative (M2) *Nf1*^ΔMΦ^ macrophages (**Figure 4A – F**). Consistent with the increase in GLUT1 expression in reparative (M2) *Nf1*^ΔMΦ^ macrophages, the expression of phosphorylated-PI3K (P-PI3K) and phosphorylated-Akt2 (P-Akt2), but not phosphorylated-Akt1 (P-Akt1), are significantly higher in reparative (M2) *Nf1*^ΔMΦ^ macrophages when compared to reparative (M2) *Nf1*^fl/fl^ macrophages (**Figure 4E – F)**. To confirm this relationship, we probed the membrane fraction of all macrophage populations from both genotypes for Akt2 and identified a 4-fold increase in P-Akt2 in reparative (M2) *Nf1*^ΔMΦ^ macrophages (**Figure 4G & H**). These data suggest that loss of neurofibromin induces GLUT1 expression through Akt2 activation.

**Figure 4.**
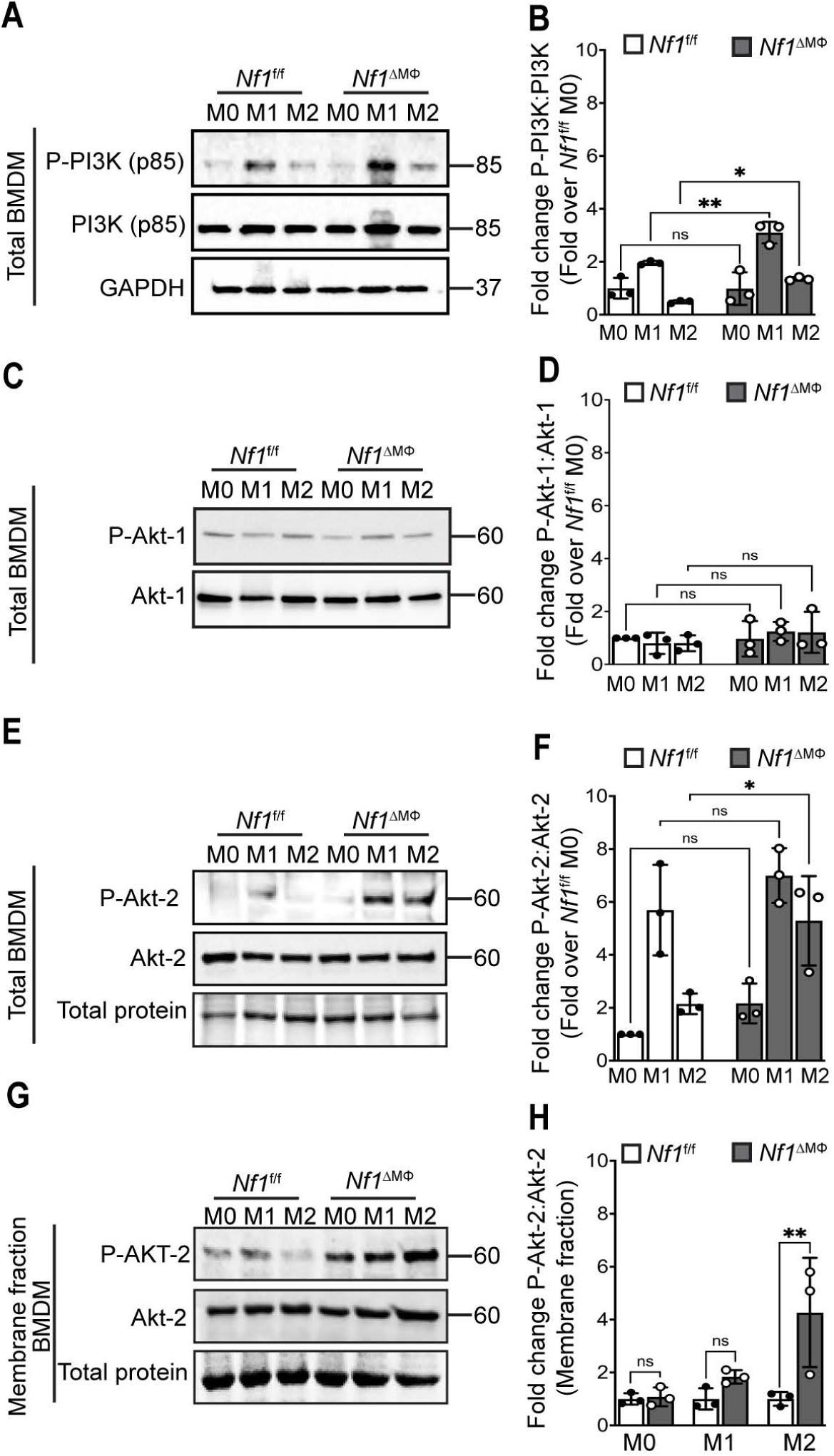
Loss of Neurofibromin in all macrophages preferentially increases Akt-2 Phosphorylation in total and in membranne fractions. Representative immunoblots and densitometry graphs showing loss of neurofibromin in macrophages increases the ratio of **(A** and **B)** P-Pl3K/Pl3K, **(C** and **D)** P-Akt-1/Akt-1, **(E** and **F)** P-Akt-2/Akt-2 expressions in total BMDM lysates and **(G** and **H)** P-Akt-2/Akt-2 expression in membrane fractions of BMDM from long bones of *Nf1*^f/f^ and *Nf1*^ΔMφ^ mice, n=3 mice per genotype, Data are expressed as mean ± SD. P values calculated using One-way ANOVA followed by Sidak’s multiple comparison text for (**B, D, F** and **H**) *P < 0 05 ** P < 0 01 *** P < 0 001 **** P < 0 0001 ns not significant).

To determine the relationship between Akt2 and GLUT1 in the absence of neurofibromin, we analyzed the membrane fraction of inflammatory (M1) and reparative (M2) *Nf1*^ΔMΦ^ and *Nf1*^fl/fl^ macrophages in the presence of CCT128930, a selective Akt2 inhibitor. Pharmacologic inhibition of Akt2 completely suppressed GLUT1 translocation to the plasma membrane of inflammatory (M1) *Nf1*^fl/fl^ macrophages with similar but diminished results seen in inflammatory (M1) *Nf1*^ΔMΦ^ macrophages (**Figure 5A – C**). In comparison, GLUT1 expression in the plasma membrane of reparative (M2) *Nf1*^ΔMΦ^ macrophages was not altered in the presence of an Akt2 inhibitor. This surprising observation led us to probe the membrane lysates of reparative (M2) macrophages from both genotypes for P-Akt1 to identify potential compensatory mechanisms resulting from pharmacologic inhibition of Akt2. In fact, we identified that Akt2 inhibition increased the expression of phosphorylated-Akt1 in both genotypes of reparative (M2) macrophages, but the increase was much more substantial in the reparative (M2) *Nf1*^ΔMΦ^ macrophages (**Figure 5D & E**). Given the relatively low expression of GLUT1 in reparative (M2) *Nf1*^fl/fl^ macrophages, we conclude the impact of Akt2 inhibition on Akt1 activation has little impact on GLUT1 localization in the presence of neurofibromin. Conversely, the loss of neurofibromin accentuates Akt1 signaling as a compensatory pathway when Akt2 is inhibited to maintain the high expression of active GLUT1 in the plasma membrane.

**Figure 5.**
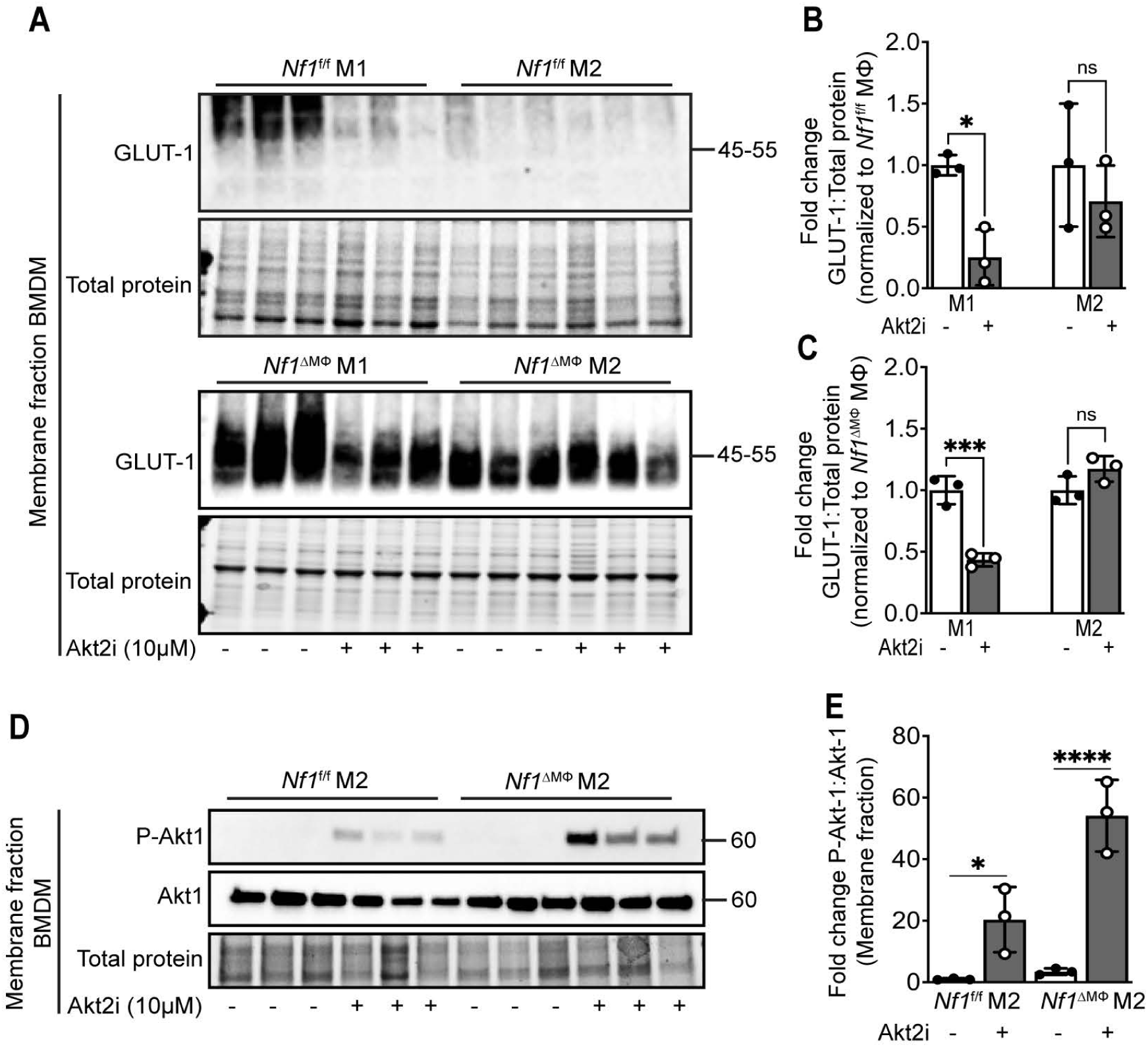
Loss of neurofibromin in macrophages promotes GLUT1 translocation to cell membrane via Akt2. Representative immunoblots and densitometry graphs showing **(A, B** and **C)** GLUT1 expression in membrane fractions of both inflammatory (M1) and reparative macrophages (M2), **(D** and **E)** P-Akt-1 and total Akt-1 expressions in membrane fractions of only M2 macrophages, from long bones of *Nf1*^f/f^ and *Nf1*^ΔMφ^ mice after Akt2 inhibitor (CCT128930, Akt2i) treatment at a dose of 10µM for 6 hours, (n=3 per genotype). Data are expressed as mean ± SD. **P** values calculated using One-way ANOVA followed by Sidak’s multiple comparison test (*P < 0.05, ** P < 0.01, *** **P** < 0.001, **** **P** < 0.0001, ns, not significant).

### Neurofibromin complexes with GLUT1 and conserved regions of Akt2

Given the compensatory effects of Akt1 in the presence of an Akt2 inhibitor, we examined the interaction between GLUT1 and neurofibromin or Akt2 by using the crystal structures of neurofibromin, GLUT1, and Akt2 to perform protein-protein docking, molecular dynamics simulation, and identify the docking score and binding affinity between molecules. **Figure 6A** shows the cartoon representation of neurofibromin docked with GLUT1 and **Figure 6B** shows GLUT1 docked with Akt2 with docking scores of −429.60 and −301.96, respectively. The binding of GLUT1 with neurofibromin is stronger than its binding with Akt2 based on their respective docking scores, indicating a more plausible binding model for neurofibromin and GLUT1 when compared with GLUT1 and Akt2. Although GLUT1 docking with neurofibromin is adjacent to the Ras-GAP domain (residues 1235-1451), few amino acid residues in this domain actively interact with residues of GLUT1 (**Supplementary Table 1**). GLUT1 binds to Akt2 via the protein kinase and AGC-kinase C terminal domain that may be independent of its binding to neurofibromin since only four amino residues i.e., ARG89, PHE90, MET98 and LEU101 (highlighted in **Supplementary Table 1**) of GLUT1 are shared between the two complexes. Two residues in GLUT1 (ARG89 & PHE90) actively interact with both Akt2 domains. Given the more plausible binding model for neurofibromin and GLUT1, Akt2 has a lower propensity to complex with GLUT1 in the presence of neurofibromin. Conversely, the absence of neurofibromin would increase the propensity for GLUT1 to interact with Akt2.

**Figure 6.**
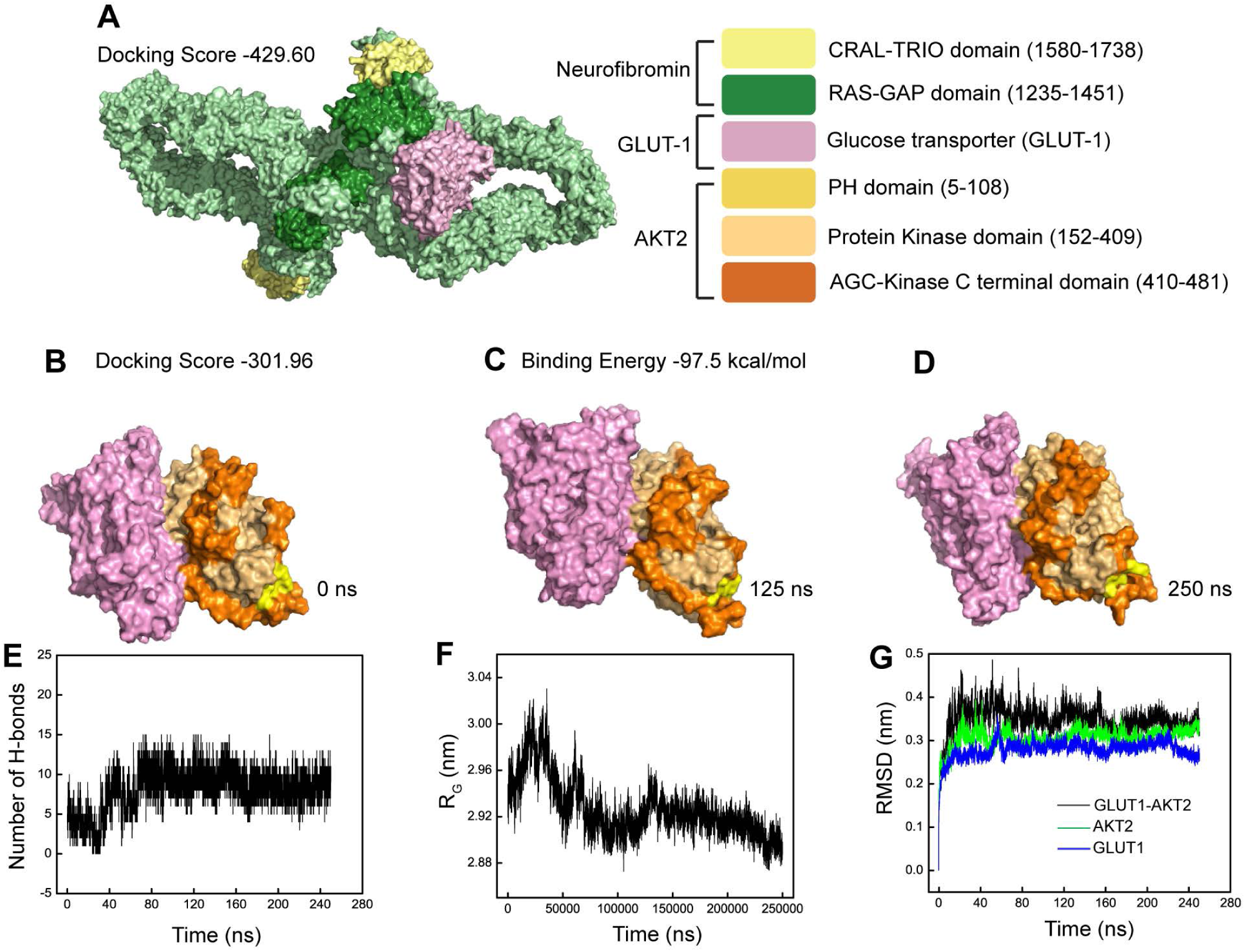
Neurofibromin complexes with GLUT1 and conserved regions of Akt2. Molecular docking and atomistic simulations implicates the complex formation of NF1-GLUT1 and GLUTI-AKT2. Surface representation of (A) NF1-GLUT1 complex having docking score of −429.60 obtained by H-dock, (B) GLUT1-AKT2 at O ns with docking score of −301.96 (C) GLUT1-AKT2 at 125 ns (D) GLUT1-AKT2 at 250 ns with binding energy of −97.5 kcal/mol obtained by simulations, plot showing (E) number of H-bonds present in GLUT1-AKT2 complex over a period of 250 ns (F) Radius of gyration, RG, of GLUT1-AKT2 complex over a period of 250 ns (G) Root mean square deviation (RMSD) of GLUT1, AKT2 and GLUT1-AKT2 complex.

To determine the stability of the GLUT1-Akt2 complex, atomistic simulations were generated for 250 ns with the conformations of the complex at 0 ns, 125 ns, and 250 ns shown in **Figure 6B-D**, respectively. The simulations estimate that the GLUT1-Akt2 complex is strong with binding energy of −97.50 kcal/mol which strengthened over time with an increase in H-bonds from 5 to 11 with consistent radii of gyration values lying between 2.89 and 2.92 nm (**Figure 6E &F**). Finally, we assessed the conformational drift of Akt2, GLUT1, and the GLUT1-Akt2 complex by calculating the Root-mean-square deviation (RMSD) of atomic positions. Equilibration of the GLUT1-Akt2 complex was reached at ∼160 ns in simulations and did not fluctuate (**Figure 6G**). Further, the overall RMSD or equilibrium value for the complex is higher than either Akt2 or GLUT1 suggesting a strong and stable complex in the absence of neurofibromin.

### Loss of neurofibromin in macrophages is sufficient to induce retinal neovascularization

Characteristic retinal neovascular lesions are observed in ∼ 80% of persons with NF1 and are reproduced in mice harboring germline *Nf1* mutations(22). Given the metabolic phenotype observed in persons with NF1 and evidence of chronic inflammation in this population, we sought to identify whether loss of neurofibromin in macrophages and subsequent inflammatory phenotypic switch might induce retinal neovascularization. Initially, we subjected WT and *Nf1* heterozygous (*Nf1*^+/−^) mice to oxygen induced retinopathy (OIR, **Figure 7A**) to demonstrate that neurofibromin expression is reduced in WT retina lysates and that F4/80 expression, a putative macrophage marker, is increased in *Nf1*^+/−^ retinas subjected to OIR (**Figure 7B – E**). We also demonstrate that *Nf1*^+/−^ retinas exhibit substantially more GLUT1 when compared with littermate WT retinas subjected to OIR (**Figure 7F & G**). Next, we subjected *Nf1*^ΔMΦ^ and *Nf1*^fl/fl^ mice to OIR and identified a significant increase in neovascular area in *Nf1*^ΔMΦ^ retinas when compared with *Nf1*^fl/fl^ retinas subjected to OIR suggesting that loss of neurofibromin in macrophages is sufficient to induce retinal neovascularization and recapitulates a feature of human NF1 (**Figure 7I & J**).

**Figure 7.**
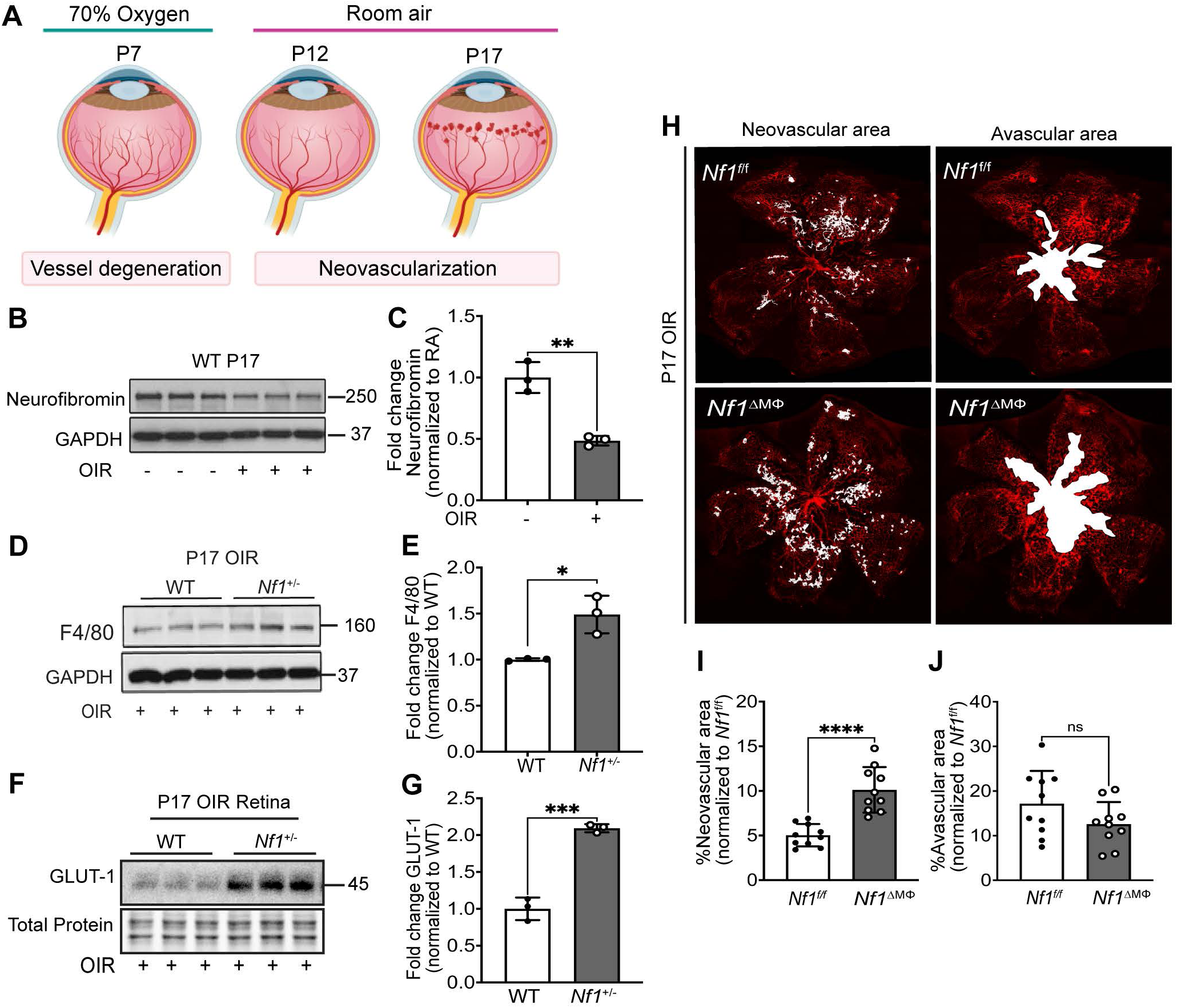
Neurofibromin is sensitive to hypoxia and loss of neurofibromin in macrophages/microglia is sufficient to induce pathological retinal neovascularization. **(A)** Schematic of the mouse Oxygen-induced Retinopathy (OIR) model. Representative immunoblots and quantifications showing **(B-C)** neurofibromin expression in P17 retinas from WT mice exposed to room air(-) or OIR (+) (n=3 mice per group), **(D-E)** F4/80 expression in P17 retinas from WT and *Nf1*^+/−^ mice exposed to OIR (n=3 mice per genotype), **(F-G)** upregulation of GLUT1 expression in P17 retinas from WT and *Nf1*^+/−^ mice exposed to OIR (n=3 mice per genotype), **(H)** Representative retinal whole mounts from P17 OIR retinas of *Nf1*^f/f^ and *Nf1*^ΔMφ^ mice stained with isolectin-B4 (red) with highlighted neovascular areas and avascular areas (white). **(I-J)** Quantification of neovascular areas and avascular area in OIR retinas were expressed as percentage of total retinal areas (n =10 retinas). Data are expressed as mean ± SD. P values calculated using 2-tailed Student’s t-test (*P < 0.05, ** P < 0.01 *** P < 0.001 **** P < 0.0001, ns, not significant).

Next, we used flow cytometry and immunohistochemistry to identify inflammatory and reparative macrophages within *Nf1*^ΔMΦ^ and *Nf1*^fl/fl^ P17 retinas subjected to OIR using the putative markers NOS2 (i.e., inducible nitric oxide synthase or iNOS) to identify inflammatory (M1) macrophages and Arginase-1 to identify reparative (M2) macrophages. We identified that NOS2-expressing retinal macrophages/microglia were increased 40-fold in P17 OIR *Nf1*^ΔMΦ^ retinas when compared with *Nf1*^fl/fl^ retinas (**Figure 8A**). Retinal macrophages/microglia expressing Arginase-1 were significantly reduced in P17 OIR *Nf1*^ΔMΦ^ retinas when compared with P17 OIR *Nf1*^fl/fl^ retinas (**Figure 8B**). Although Arginase-1 expressing macrophage/microglia were rarely identified in the retinas from either genotype subjected to OIR, the number of CD11b^+^/F4/80^+^ retinal macrophages/microglia that co-express NOS2 and Arginase-1 was increased ∼40-fold in P17 OIR *Nf1*^ΔMΦ^ retinas when compared with P17 OIR *Nf1*^fl/fl^ retinas (**Figure 8C & S3**). Finally, we examined neovascular tufts from *Nf1*^ΔMΦ^ and *Nf1*^fl/fl^ P17 retinas subjected to OIR to identify inflammatory and reparative peri-vascular macrophages. Similarly, we identified that inflammatory CD86-expresssing macrophages were more common in the neovascular tufts of *Nf1*^ΔMΦ^ retinas when compared with *Nf1*^fl/fl^ retinas subjected to OIR, but no difference in reparative CD206 macrophage content was identified in P17 OIR retinas from both genotypes (**Figure 8D – G**). Taken together, loss of neurofibromin in macrophages facilitates the accumulation of perivascular inflammatory (M1) macrophages to promote retinal neovascularization.

**Figure 8.**
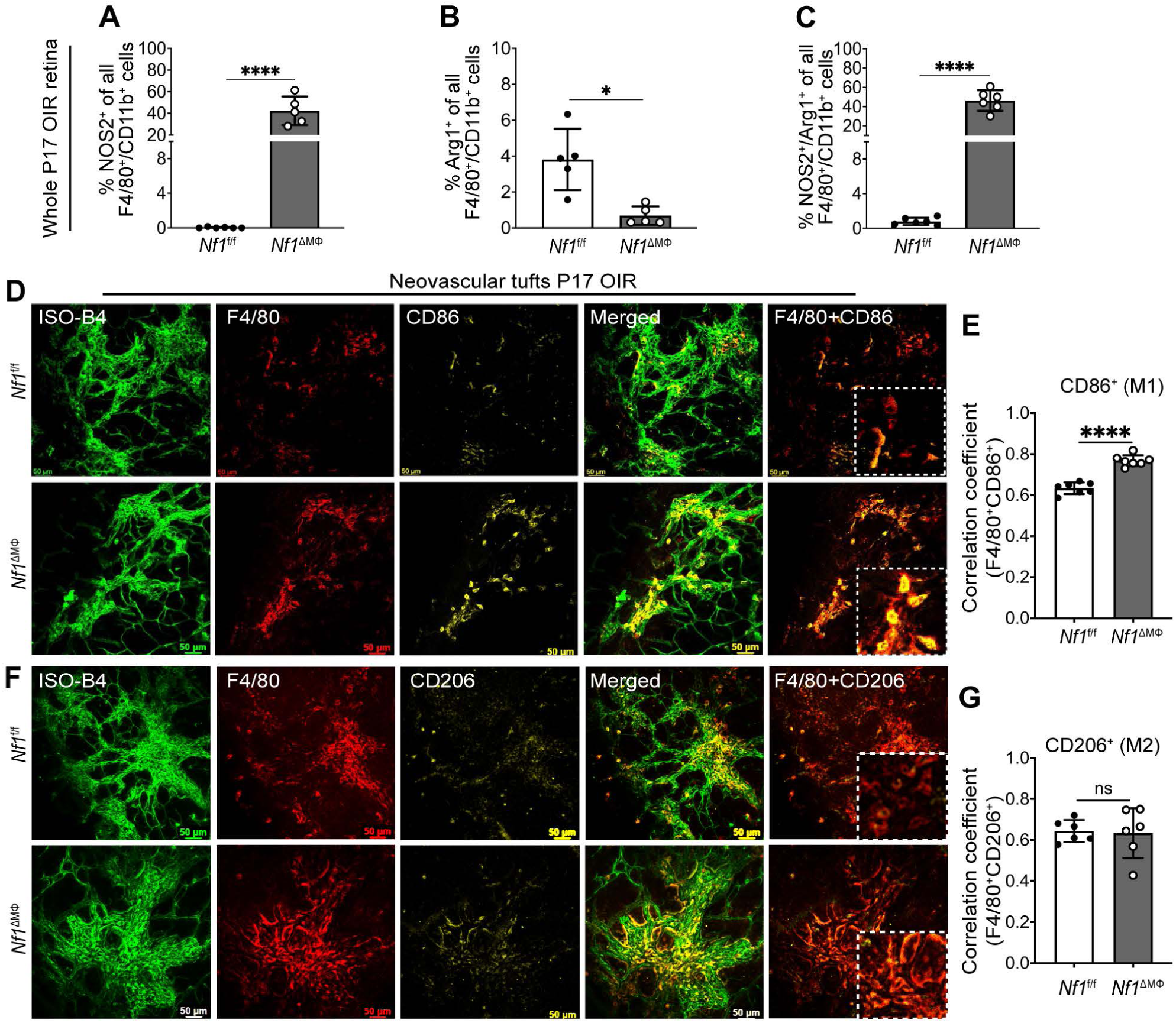
Loss of Neurofibromin Promotes Inflammatory Macrophage/Microglia infiltration and localization to sites of Neovascular Tufts in the retina. Flow cytometric analysis of whole P17 retinas from *Nf1*^f/f^ and *Nf1*^ΔMφ^ mice exposed to OIR expressing macrophage or microglial markers. Plots represent percentage of viable **(A)** inflammatory macrophage/microglia (F4/80^+^/CD11b^+^/NOS2^+^), **(B)** reparative macrophage/microglia (F4/80^+^/CD11b^+^/Arg1^+^), and **(C)** macrophage/microglia with mixed phenotype (F4/80^+^/CD11b^+^/NOS2^+^/Arg1^+^) in P17 OIR retinas (n=5-6 mice per genotype; total of 10-12 retinas in each genotype). Confocal images of retinal flat mounts at P17 OIR from *Nf1*^f/f^ and *Nf1*^ΔMφ^ mice, stained with isolectin-B4 (vessel stain}, F4/80 and either **(D)** CD86 (inflammatory macrophage/microglia) or **(F)** CD206 (reparative macrophage/microglia) to show the localization of macrophage/microglia near neovascular tufts. Pearson’s correlation coefficient for F4/80 with either CD86 **(E)** or with CD206 **(G)** to depict colocalization (n = 6 retinas in each group). Scale bars: 50um **(D** and **F);** 10um (inset). Data are expressed as mean ± SD. P values calculated using 2-tailed Student’s t-test (*P < 0.05, ** P < 0.01, *** P < 0.001, **** P < 0.0001, ns, not significant).

The observation that retinal macrophages/microglia from *Nf1*^ΔMΦ^ mice subjected to OIR are more likely to exhibit features of an inflammatory phenotype and co-express putative markers of inflammatory (M1) and reparative (M2) macrophages, we probed bone-marrow derived *Nf1*^ΔMΦ^ and *Nf1*^fl/fl^ macrophages for inflammatory cytokines, NOS2 and NF-κB expression. In **Figure 9A-C**, we identified that both inflammatory (M1) and reparative (M2) *Nf1*^ΔMΦ^ macrophages express much higher quantities of IL-6, IL-1β, and MCP-1, which are cytokines that have been linked to NF1 in either human serum or murine models. We have also observed increased level of NOS2 in all *Nf1*^ΔMΦ^ macrophages, but the increase was most dramatic (10-fold) in reparative (M2) macrophages. Taken together, the loss of neurofibromin polarizes macrophages toward a pro-inflammatory phenotype irrespective of the polarizing agents.

**Figure 9.**
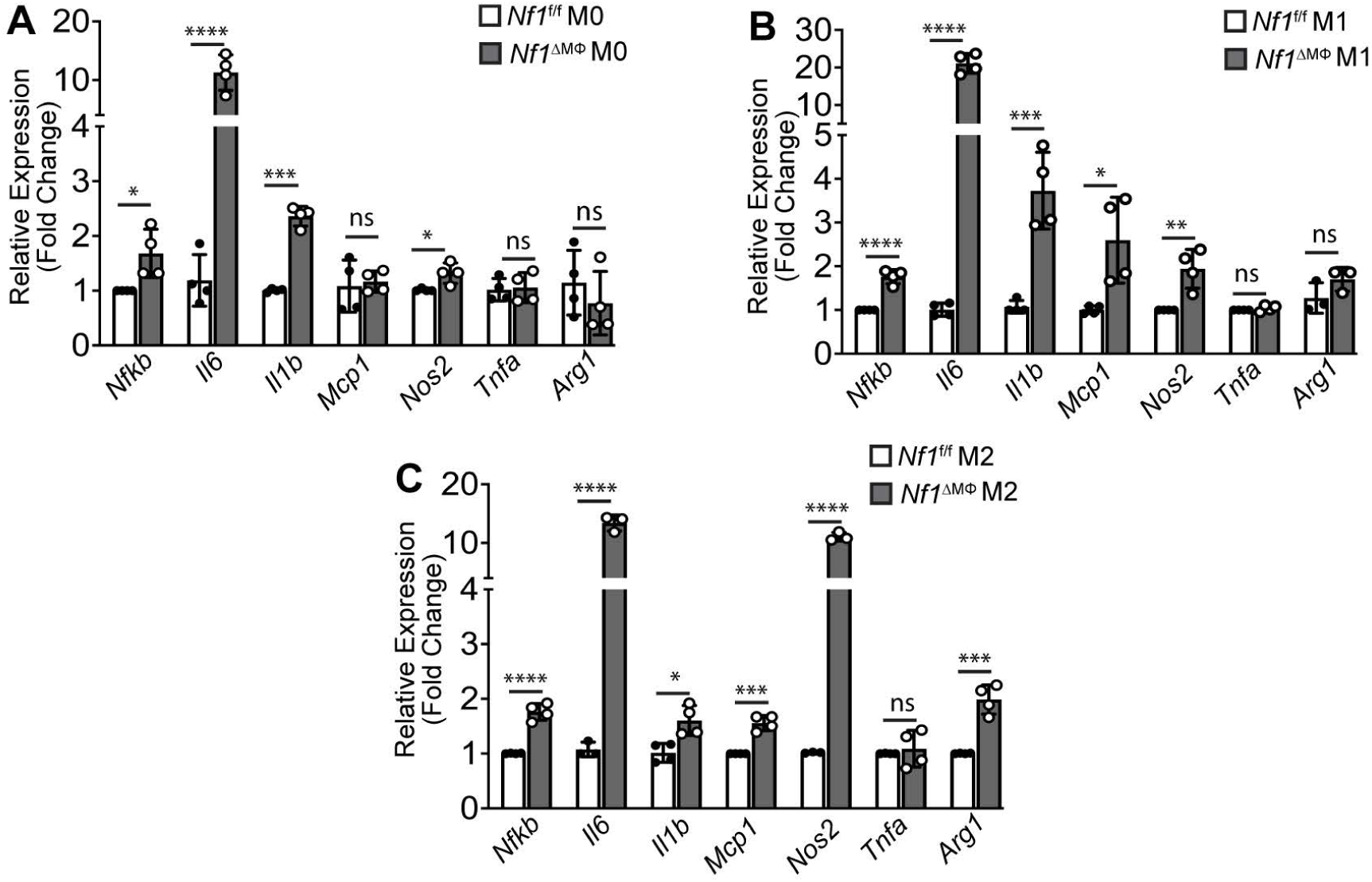
Loss of neurofibromin results elevated relative mRNA expression of inflammatory cytokines, NF-kB and NOS2 and polarizes macrophages to mixed phenotypes. **(A, B** and **C)** qRT-PCR analysis of relative mRNA expressions of inflammatory cytokines, including IL-6, IL-1β, MCP-1, TNF-α, NOS2 and NF-kB in **(A)** control (M0), **(B)** inflammatory (M1), and **(C)** reparative (M2) macrophages derived from long bones of *Nf1*^f/f^ and *Nf1*^ΔMφ^ mice. RPLP0 is considered as endogenous control in this study (n=4 mice per genotype, each data point represents the mean of two technical replicates of each mice). Data are expressed as mean ± SD. P values calculated using 2-tailed Student’s t-test (*P < 0.05, ** P < 0.01, *** P < 0.001, **** P < 0.0001, ns, not significant).

### GLUT1 and phosphorylated-Akt2 co-localize within macrophages from NF1-associated tumors

It is well documented that macrophages are prevalent in neurofibromas and malignant peripheral nerve sheath tumors (MPNST) and correlate with disease progression (23–25). However, the reason why macrophage density increases with disease progression is still not well explained. Therefore, we finally sought to confirm our observations from lineage-restricted *Nf1*-mutant mice in relevant human tissue. Procuring human retina tissue from persons with NF1 is not practical; therefore, we examined tumor samples from persons with NF1 for expression of GLUT1 and P-Akt2 within perivascular macrophages. We identified 5 plexiform neurofibromas (pNF) and MPNST samples from 4 patients (mean age = 31 years, 50% male; details in **Supplementary Table 2**) and probed paraffin embedded sections for GLUT1 and P-Akt2. Both GLUT1 and P-Akt2 co-localized with 80% of the CD68-expressing macrophages within these tumor samples suggesting a high degree of GLUT1 activation in pNF and MPNST-associated macrophages (**Figure 10A & B**).

**Figure 10.**
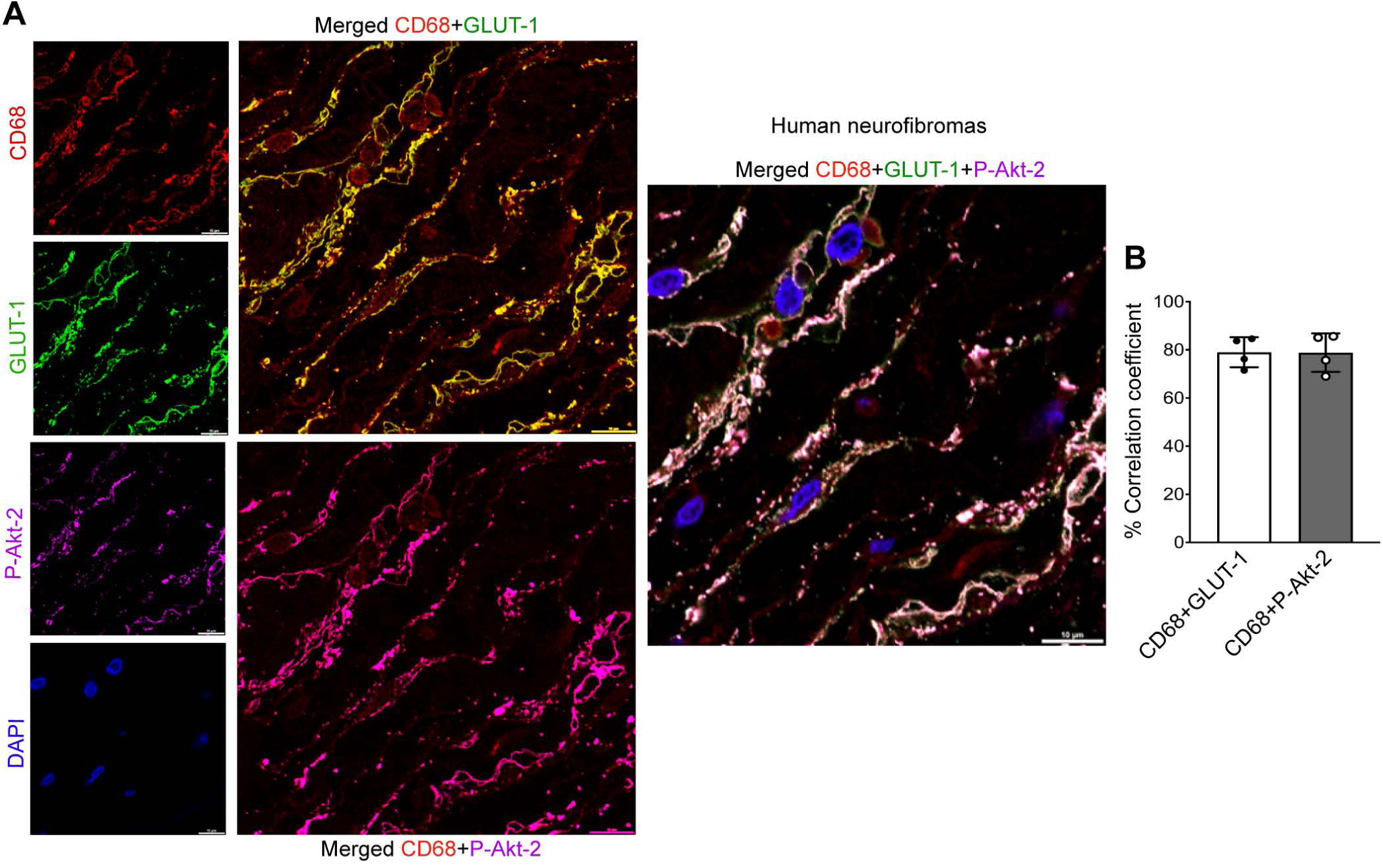
Loss of neurofibromin results GLUT1 and Akt2 co-localization within CD6S^+^ macrophages from NF1-associated tumors. (A) Representative high magnification confocal images of NF1-associated tumor cross-section from human patients showing colocalization of GLUT1 and P-Akt2 with CD68^+^ macrophages, Scale bars: 10um. (B) Percent Pearson’s correlation coefficient for CD68 with either GLUT1 or P-Akt2 to depict colocalization (n = 4 patients with NF1). Each data point reflects the values to each patient.

## Discussion

Neurofibromatosis type 1 is a multi-system disease resulting from dominantly inherited mutations in the *NF1* tumor suppressor gene, encoding neurofibromin. Despite the relative inconsistency in the timing and penetrance of any one manifestation in persons with NF1, human cohort studies have identified a characteristic metabolic and anthropometric phenotype characterized by short stature and reduced adiposity as well as a strong protection from insulin resistance and diabetes. A recent registry study of nearly 2,500 persons with NF1 demonstrated that persons with NF1 have statistically lower odds of hospitalization for endocrine-related diseases when compared with the general population(26). Of the 11 main diagnostic categories for hospitalization days, only endocrine-related diagnoses produced a negative association with NF1. Interestingly, the risk for endocrine-related hospitalization was reduced by 80 and 40% in males and females with NF1, respectively. When examining individual diagnoses within this category, the prevalence of diabetes mellitus type 1 was reduced 2.5-fold while diabetes mellitus type 2 prevenance was similar to the general population. A more recent Finnish registry study explored this potential relationship with a specific focus on diabetes mellitus(7). In comparison to non-NF1 siblings and non-related matched controls, the hazard ratio for diabetes mellitus type 1 in the NF1 cohort was 0.55 and 0.58, respectively. Again, males with NF1 seem to display greater protection from diabetes mellitus type 1 when compared with male siblings and male controls with hazard ratios of 0.26 and 0.29, respectively. Finally, those persons with NF1 who were diagnosed with diabetes mellitus type 1 appear to require less insulin when compared with sibling and non-sibling controls. Experimental human studies identified a nearly 90% lower risk of persons with NF1 having a fasting blood glucose level above 100 mg/dL when compared with matched non-NF1 controls, but no group differences in insulin or blood glucose values after administration of a dextrose bolus(5, 6). We have demonstrated similar findings in *Nf1* heterozygous mice(8). Taken together, these human registry data and experimental studies are highly suggestive that mutations in the *NF1* gene harbor a hormone-independent relationship to enhanced glucose internalization and/or metabolism, which serves as the theoretical basis for our experiments.

Our long-standing interest is understanding why persons with NF1 experience chronic inflammation and the untoward effects of acute and sustained inflammation on this population. Macrophages are a rational population for mechanistic studies in NF1 and are pathogenically implicated in multiple models of NF1. Human and murine NF1/Nf1 tumors are replete with macrophages and macrophage density correlates with tumor growth and malignant transformation(15, 23, 25, 27). Descriptive studies have consistently pointed to an inflammatory signature in these tumors with a preponderance of so-called inflammatory macrophages and high expression of inflammatory cytokines corresponding with a lower-than-expected number of so-called reparative macrophages. The 10 to 20-fold increase in secreted IL-6 from neurofibromin-deficient macrophages is of particular interest as several publications have demonstrated higher plasma IL-6 concentrations in persons with NF1 when compared with matched controls(13, 14, 28).

Here, we converge these observations to demonstrate that loss of neurofibromin does impart a rapid and sustained entrainment of glucose into macrophages that is mediated through GLUT1. We also demonstrate that the relationship between GLUT1 and neurofibromin is complex within these cells such that the increase in membrane-bound GLUT1 in neurofibromin-deficient cells is not limited to so-called inflammatory (M1) macrophages, which are highly reliant on GLUT1 to internalize glucose. Rather, we identified that so-called reparative (M2) macrophages also exhibit high membrane bound GLUT1 in the absence of neurofibromin corresponding with a reduction in cytoplasm GLUT1 content. This imbalance in functional GLUT1 availability may skew the so-called reparative macrophages to exhibit features that are more commonly identified in inflammatory macrophages. In keeping with this line of thinking, both inflammatory and reparative neurofibromin-deficient macrophages express much higher inflammatory cytokine content when compared with wildtype macrophages. Further, we demonstrate that loss of neurofibromin in macrophages alone is sufficient to induce retinal neovascularization and that the neovascular tufts are invested with inflammatory macrophages. Again, we find that retinal macrophages in *Nf1*^ΔMΦ^ retinas subjected to OIR are not dichotomous with some macrophages expressing putative markers of both macrophage subpopulations. While this same phenomenon is seen in control retinas, it is much less pronounced and further supports our hypothesis that loss of neurofibromin promotes both classic inflammatory macrophage function and phenotypic switching of classic reparative macrophages.

The ability of neurofibromin-deficient macrophages to take on properties of both inflammatory and reparative macrophages provides insight into many manifestations of NF1 that have been linked to inflammatory changes, cells, or cytokines in animal models or human tissue. It has been speculated that macrophages are part of a secondary response to a disease-initiating event and propagate pathology. For example, multiple investigative teams have demonstrated that inflammatory macrophages are abundant in NF1 tumors and that primary tumor cells secrete growth factors and cytokines that recruit both wildtype and neurofibromin-deficient macrophages toward the stimulus(16, 25, 29). In line with this paradigm, we have demonstrated that the increased sensitivity to growth factors and cytokines observed in neurofibromin-deficient macrophages may extend beyond a mere signaling amplification to include an increased propensity to sample their extracellular environment through endocytosis(10, 30, 31). However, this line of thinking does not elucidate the fundamental question of why inflammatory macrophages are abundant in tumors and other manifestations of NF1. Here, we identify a conserved feature of neurofibromin-deficient macrophages and tissue that enhanced glucose internalization via a constitutively active GLUT1 underlies these previously recognized features. Given this defining feature of inflammatory macrophages and experimental evidence suggesting their role in driving NF1-related pathologies such as tumor growth and malignant transformation, targeting glucose internalization or glycolysis specifically in macrophages is an attractive therapeutic approach for persons with NF1.

Targeting GLUT1 directly is not a simple prospect owing to its necessity in maintaining basal glucose supply to most cells. We demonstrate that membrane bound GLUT1 is controlled by Akt2, but loss of neurofibromin complicates this relationship particularly in so-called reparative (M2) macrophages. In the presence of neurofibromin, Akt2 binding to GLUT1 is impaired and likely explains the clear loss of membrane bound GLUT1 following Akt2 inhibition. When neurofibromin is absent, GLUT1 is much more likely to bind to Akt2, which in turn enhances GLUT1 expression in the membrane and explains the increase in glucose internalization to drive inflammatory macrophage phenotypic switch. Further, Akt2 inhibition becomes complicated in that Akt1 appears to take on some of the putative Akt2 signaling in the absence of neurofibromin. These data imply that targeting the Akt2-GLUT1 pathway for persons with NF1 needs to be macrophage-specific and incorporate combination therapies targeting PI3K and/or both Akt isoforms.

Finally, we provide compelling animal and human evidence that macrophage-mediated inflammation is critical to NF1-related angiogenesis. First, neurofibromin is sensitive to hypoxia. Following exposure to hyperoxia from P7 to P12, animals returned to normoxia experience a relative hypoxia that drives vascularization of the retina with peak neovascularization occurring at P17. We demonstrate that retinal neurofibromin expression is suppressed during the hypoxia phase of OIR and this corresponds with an increase in retina macrophage density and specifically increases inflammatory macrophage content within the retinal neovascular tuft. Although persons with NF1 have characteristic neovascularization of the retina and choroid that are now part of the diagnostic criteria for NF1(22, 32–34), the implications of these findings extend beyond the retina since perivascular cuffing is a common feature of NF1 tumors. Further, we demonstrate that the relationship between over-expression of GLUT1 and loss of neurofibromin is conserved at the tissue level since GLUT1 expression is increased nearly 2-fold in *Nf1*+/− retinas when compared with WT retinas despite macrophages representing a small fraction of the cell pool within the retina. Similar results are observed in human NF1 tumors with high concordance of GLUT1 and Akt2 within tumor macrophages.

In conclusion, we demonstrate that loss of neurofibromin facilitates GLUT1 translocation to the membrane of macrophages to facilitate a phenotypic switch between classically reparative and inflammatory macrophages. This relationship is enhanced via Akt2, which is more easily bound to GLUT1 in the absence of neurofibromin and activates its glucose internalization properties. These mechanistic findings provide a rational explanation for the high numbers of inflammatory macrophages observed within NF1 tumors and supports a role for inflammatory macrophages in promoting neovascularization in the retina and tumors of persons with NF1.

## Methods

### Sex as a biological variable

Both male and female mice were used for all *in vitro* and *in vivo* studies. Tissue samples from both male and female persons with NF1 were analyzed. Sex was not considered as a biologic variable for any studies.

### Cells

Murine bone marrow-derived macrophages (BMDM), and human THP-1-derived macrophages were cultured in high glucose Dulbecco’s modified Eagle’s Medium (DMEM) medium containing 10% FBS, 100 U/mL of penicillin G and 100 mg/mL streptomycin. THP1-derived macrophages were kindly gifted from Dr. Gabor Csanyi (Augusta University, Augusta, GA). Media was supplemented with either Macrophage colony-stimulating factor (M-CSF; 20 ng/ml, PMC2044; Invitrogen) or 20% L929 cell-derived conditioned media to differentiate bone marrow-derived monocytes into macrophages. Cells were cultured in the absence of L929 cell-derived conditioned media for at least 24 hours prior to experiments. Human THP-1 monocytes were differentiated using PMA (200nM). Both BMDM and THP-1 macrophages were polarized into so-called inflammatory macrophages (M1) using IFN-ɣ (20ng/ml) and LPS (100ng/ml) and reparative macrophages (M2) using IL-4 (20ng/ml) for 16 hours in serum-free high glucose DMEM media.

### Primary bone marrow-derived macrophages

Murine bone marrow monocytes were isolated and differentiated as described by Trouplin *et. al* with modifications(35). Briefly, the femur and tibia were dissected, cleaned, and flushed with sterile PBS using a 10mL syringe and a 25-gauge needle. Bone marrow was spun down and resuspended in differentiation media (high-glucose DMEM, 10% FBS, 20% L929 cells-derived conditioned media and 1% penicillin/streptomycin). After thoroughly being pipetted to obtain a single cell suspension, the bone marrow was plated on 3×10cm petri dishes (uncoated). This was determined to be day 1. On day 3, the media was removed from the plates and replaced with fresh differentiation media. On day 7, differentiation media was replaced with polarization DMEM media containing IFN-ɣ (20ng/ml) and LPS (100ng/ml) for inflammatory (M1) and IL-4 (20 ng/ml), for reparative (M2) macrophages and were incubated for 16 hours. On day 8, the macrophages were ready to use for experiments.

### Animals

Mice were housed in the animal facility of Augusta University under a 12-hour light/dark cycle with ad libitum access to food and water. *Nf1*-heterozygous (*Nf1*^+/−^), *Nf1^flox/flox^*;LysMCre^+^ (*Nf1*^ΔMΦ^) and *Nf1^flox/flox^*;LysMCre^-^ (*Nf1*^fl/fl^) mice were obtained from Tyler Jacks, Ph.D., (Massachusetts Institute of Technology, Cambridge, MA). *Nf1^+/−^* mice were backcrossed 13 generations into the C57BL/6J strain. *Nf1*^f/f^ mice were obtained from Luis Parada (University of Texas Southwestern Medical Center, Dallas, TX, USA). *Nf1*^f/f^ and LysMCre^+^ mice were crossed till second generation to produce *Nf1*^f/f^ and *Nf1*^ΔMΦ^ mice which were used as control and myeloid cell-restricted knock out mice in experiments, respectively. Mice were genotyped by PCR as described(36). WT and *Nf1*^+/−^ mice (C57BL/6J) were crossed to generate experimental progeny. Littermate male mice, between 8 to 12 weeks of age, were used for experiments.

### Glycolysis and Metabolic Analysis

Cellular bioenergetics were measured using an XF96 analyzer (Agilent Seahorse XF Technology, Santa Clara, CA). Macrophages (THP-1, and BMDM from *Nf1*^fl/fl^ and *Nf1*^ΔMΦ^ mice) were seeded at 50,000 cells per well and 24 hours later polarizing cytokines (refer section *Primary bone marrow-derived macrophages*) were added and incubated for 16 hours. Glycolysis in polarized BMDM was measured using Seahorse XF glycolysis stress test kit (Agilent Technologies; 103020-100) as per manufacturer’s protocol. After one hour of equilibration of BMDM in DMEM, glycolysis was measured with injections of glucose (10 mM), oligomycin (1.5 uM) and 2-DG (50 mM). The energy phenotype was measured by adding both oligomycin (1.5uM) and carbonyl cyanide 4-(trifluoromethoxy) phenylhydrazone (FCCP; 1.5uM) simultaneously and measuring the change in ECAR and OCR.

### Glucose Uptake assay

Glucose uptake was measured using a fluorescent 2-NBDG Glucose Uptake Assay Kit (BioVision; K682-50) as per manufacturer’s instruction with slight modifications. Briefly, polarized BMDM (M0, M1 and M2) derived from long bones of *Nf1*^f/f^ and *Nf1*^ΔMΦ^ mice were seeded at a density of 500,000/well in 24-well plate in high glucose DMEM with 10% FBS. After 24 h, cells were serum starved in 0.2% serum rich DMEM for 30 minutes. Starved cells were then incubated with and without BAY-876 inhibitor (Cayman Chemical Company; 19961, 50nM) in 0.2% serum-rich DMEM for 30 minutes more. After incubation of BAY-876, cells were incubated with a FITC-labeled glucose analog (2-deoxy-2-[(7-nitro-2,1,3-benzoxadiazol-4-yl) amino]-D-glucose) for 30 minutes at 37°C and fluorescence was measured at a wavelength of 488 nm with flow cytometry.

### Flow Cytometry

Cells ells were stained according to the protocol described previously by Zaidi *et.al*(37). Briefly, retinas from *Nf1*^f/f^ and *Nf1*^ΔMΦ^ P17 OIR pups were collected and single cell suspension from both retinas together was made by gentle mechanical dissociation in ice-cold staining buffer (BD Biosciences; 554656). Cells were centrifuged, pellet were collected and assessed for viability using Zombie NIR fixable viability kit (BioLegend; 423105) for 30 minutes at 4°C. Cells were then incubated in FcR (CD16/CD32) Blocking reagent (BD Biosciences; 553141) for 20 minutes at 4°C. For surface marker staining, cells were incubated with fluorescently conjugated primary antibodies, PerCP-CD11b (BioLegend; 101230) and FITC-F4/80 (BioLegend; 123108) for 30 minutes at 4°C in dark, followed by washing with ice-cold staining buffer and centrifuged at 300g for 5 minutes. For intracellular staining, cells were permeabilized and fixed with BD Cytofix/Cytoperm buffer (BD Biosciences; 555028) for 20 minutes at 4°C in dark, followed by washing with BD Perm/Wash buffer (BD Biosciences; 555028). Cells were washed, centrifuged at 300g for 5 minutes and were stained with primary antibodies, PE-Cy7-iNOS (Invitrogen; 25-5920-82) and PE-Arg-1 (Invitrogen; 12-3697-82) for 30 minutes at 4°C in dark. Finally, cells suspension was washed, centrifuged and filtered in 5ml tube with cell strainer cap (40um) and analyzed on Acea NovoCyte Penteon flow cytometer (Agilent). Novocyte software was used to analyze the data. Unstained and Fluorescence minus one (FMO) controls were used for gating. Compensation was done using UltraComp eBeads plus compensation beads (Invitrogen; 01-3333-42).

### Protein extraction and Western blot

Whole protein lysates were prepared from single cell suspension of BMDM from *Nf1*^f/f^ and *Nf1*^ΔMΦ^ mice or cell lines or tissue with 1X Pierce RIPA buffer (ThermoFisher Scientific, 89900) supplemented with 1% proteinase/phosphatase inhibitor cocktail following established protocol(38, 39). Protein concentration was determined by Pierce bicinchoninic acid (BCA) protein assay (ThermoFisher Scientific, 23227) according to the manufacturer’s instructions. Equal amount of protein (20mg) was loaded on 10-12% TGX Stain-free SDS-PAGE gel (Bio Rad, 1610183 and 1610185) and transferred to PVDF membrane (Bio-Rad, 1620177). Primary antibodies used in this study were as follows: Rabbit GLUT-1 (abcam; ab115730, 1:100,000), Rabbit Na/K ATPase (abcam; ab76020, 1:10,000), Neurofibromin (Bethyl 1:1000), Mouse GAPDH (Santa Cruz; sc-365062), Rabbit phospho-PI3K p85 (CST; 4228S, 1:1000), Rabbit PI3K (CST; 4292, 1:1000), Rabbit phospho-Akt-1 (Ser473) (CST; 9018S, 1:1000), Rabbit Akt-1 (CST; 2938T, 1:1000), Rabbit phospho-Akt-2 (Ser474) (CST; 8599S, 1:1000), Rabbit Akt-2 (CST; 3063T, 1:1000), Rat F4/80 (Bio-Rad, MCA497R),). Membranes were incubated with HRP-conjugated secondary antibodies for 1 hour at room temperature. Membranes were developed using Pierce ECL western blotting substrate (ThermoFisher Scientific, USA). Images were taken with the ChemiDoc MP system (Bio-Rad), and band densities were quantified in triplicates using Image Lab software (Bio-Rad). Data were normalized either by total protein or GAPDH as loading control.

### RNA isolation, cDNA synthesis and Quantitative Real-time PCR (qRT-PCR)

Total RNA from polarized BMDM (M0, M1, M2) from *Nf1*^f/f^ and *Nf1*^ΔMΦ^ mice were extracted using RNeasy Mini kit (Qiagen; 74104). The resultant sample of total RNA (500ng-1μg) was utilized as a template for complementary DNA (cDNA) synthesis using either iScript^TM^ cDNA synthesis kit (Bio-Rad; 1708891) or high-capacity M-MLV reverse transcriptase (Invitrogen). qRT-PCR was performed using cDNA (diluted 1 in 10 with nuclease free water), on a StepOne Plus System (Applied Biosystems) or ABI 7500 Real-Time PCR System (Applied Biosystems, Foster City, CA) (Applied Biosystems, Foster City, CA, US), using iTaq Universal SYBR Green Supermix (Bio-Rad, 1725121) with the gene-specific primers(40, 41). The primers used in this study were either obtained from TaqMan gene expression assay (Applied Biosystems) or Integrated DNA Technologies (Coralville, IA, US). Quantification of relative gene expression was calculated with the efficiency-corrected 2^-ΔΔCT^ method using murine GAPDH or RPLP0 as the internal controls for normalization following Livak *et. al* method(42) and data were presented as fold change relative to control groups. Details of primer sequences used for genes are provided in supplemental information.

### Subcellular Protein Fractionation of BMDM

To determine the facilitative GLUT1 translocation and P-Akt-2 expression in membrane and cytoplasm, subcellular fractionation of polarized BMDM from *Nf1*^f/f^ and *Nf1*^ΔMΦ^ mice was performed using Subcellular protein fractionation kit for cultured cells (ThermoFisher Scientific; 78840) according to manufacturer’s protocol with slight modifications. Briefly, BMDM at a density of 10 million for each condition (M0, M1 and M2) were seeded in 10 cm plates, differentiated for 7 days using 20% L929 cells-derived condition media, and polarized using specific cytokines into so called M1 and M2 macrophages (refer section *Isolation of mouse macrophages, differentiation and polarization*). Cytoplasmic fractions were isolated first by incubating cells in cytoplasmic extraction buffer for 15 minutes at 4°C, followed by collection of membrane fractions using membrane extraction buffer for 15 minutes at 4°C. Both fractions were used later for immunoblotting.

### Co-immunoprecipitation

To determine the complex formation between NF1 and GLUT1 in BMDM from *Nf1*^f/f^ mice, co-immunoprecipitation was performed using immunoprecipitation kit (ThermoFisher Scientific; 10007D) according to manufacturer’s protocol with slight modifications. Briefly, the BMDM from *Nf1*^f/f^ mice were isolated, differentiated and polarized to M1 and M2 macrophages (refer section *Primary bone marrow-derived macrophages*). Cells were subsequently washed once with PBS, followed by lysis in 2X RIPA buffer with 1% proteinase/phosphatase inhibitor cocktail to make lysates. Firstly, 50μl of Dynabeads were incubated with 8ug of GLUT1 antibody (CST, 12939) for 2 hours at 4°C on rotation. The Dynabeads-antibody complex made was clarified using DynaMag™-2 Magnet (ThermoFisher Scientific; 12321D), followed by washing in antibody binding and washing buffer by gentle pipetting. This Dynabead-antibody complex was incubated with lysates (800ug of protein) overnight at 4°C on rotation to allow the antigen to bind the Dynabeads-antibody complex. Next day, Antigen-Dynabeads-antibody complex was washed in washing buffer and separated. The Antigen-Dynabeads-antibody complex was resuspended in 2X sample buffer and heated for 10 minutes at 70°C. The complex was loaded onto the gel for immunoblotting. Secondary HRP-conjugated antibody or TidyBlot HRP-conjugated (Bio-Rad; STAR209PA, 1:400) were used to develop the bands.

### Protein-Protein Docking

The X-ray crystal structures of glucose transporter 1 (GLUT1: PDB ID code 4PYP), AKT2 (PDB ID code 8Q61), and neurofibromin 1 (NF1:PDB ID code 8K4L) obtained from the Protein Data Bank (PDB). Prior to docking, all heteroatoms present in the structures were removed. The modeling process was performed using the HDOCK server (http://hdock.phys.hust.edu.cn/) which employs computational algorithms to predict the three-dimensional structures of protein complexes based on known interactions and structural data. The knowledge-based iterative scoring function, ITScorePP or ITScorePR, determines the docking scores in Hdock. Since the docking score has not been calibrated to the experimental data, it should not be interpreted as the genuine binding affinity between two molecules; rather, a higher negative docking score indicates a more plausible binding model.

### Atomistic simulations

Molecular dynamics simulations were conducted using Gromacs package 2018.1. Initially the protein-protein complex of GLUT1-AKT2 was setup with amber99sb-ILDN forcefield in a cubic box whose sides were kept 1nm apart from complex. The model was solvated using TIP3P water molecules and subjected to energy minimization through 5,000 steps of the steepest-descent method. To neutralize the systems, counterions were added. Subsequently, a second round of energy minimization was conducted with an additional 5,000 steps of the steepest-descent method. Two rounds of equilibration were performed: firstly, a 0.1 nanosecond equilibration at constant number, volume, and temperature using a time step of 0.002 nanoseconds, employing the V-rescale algorithm for temperature coupling at 310 Kelvin with a coupling time constant of 0.5 picoseconds, separately applied to proteins/complexes, and solvent; secondly, a 1 nanosecond equilibration at constant number, pressure, and volume with a time step of 0.002 nanoseconds, utilizing Nose–Hoover temperature coupling and Parrinello–Rahman pressure coupling. The system’s pressure was semi-isotropically coupled using the Berendsen algorithm at 1 bar, with a compressibility of 4.5 × 10^-5^ and a coupling time constant of 5.0 picoseconds. The production run was conducted for 250 nanoseconds without any restraints, employing a time step of 0.002 nanoseconds. A neighbor-list generation and Coulomb and Lennard-Jones interactions cutoff of 1.2 nanometers were selected, with Particle–Mesh–Ewald summation employed for electrostatic interactions. End-state free energy calculations with GROMACS were conducted by using a tool gmx_MMPBSA(43).

### Oxygen-Induced Retinopathy

To induce retinal pathological angiogenesis in *Nf1*^f/f^ and *Nf1*^ΔMΦ^ mice, the OIR model was used, as previously described(44). Nursing dams with post-natal day 7 (P7) pups were housed in a sealed chamber and exposed continuously to 70% oxygen. At P12, the animals were returned to room air (21%). Pups with body weight less than 4 g at P17 were excluded from analysis. Retinas from P17 pups were collected for experiments. Data from six pups per litter were used for all experiments.

### Retina dissection and Immunofluorescence staining

For quantification of retinal neovascularization in retina whole-mount at P17 OIR stage, eyes from pups exposed to oxygen were enucleated and fixed in 4% PFA/PBS overnight at 4°C. The retinas were dissected out, washed in PBS for 10 minutes in a shaker. Retinas were then permeabilized and blocked in blocking solution (10% donkey serum and 1% Triton X-100 in PBS) for 1 hour at room temperature. After washing in PBS, retinas were stained with Alexa Fluor 594-conjugated Isolectin GS-IB_4_ (Invitrogen; I21413, 1:100 dilution in 3% donkey serum with 0.3% Triton X-100) overnight at 4°C. Next day after washing three times in PBS, the retinas were mounted on glass slides using mounting media without DAPI (Vector Laboratories; H-1000). Images were acquired at 20X magnification with Leica Stellaris confocal microscope. Quantification of avascular area and neovascular tufts was performed in Adobe Photoshop, as previously described by Connor *et. al*(45).

For whole-mount immunofluorescence staining for macrophage/microglia at P17 OIR stage, eyes were processed, fixed and permeabilized similarly as for Isolectin-B4 staining. Fixed retinas were then stained with Alexa Flour-conjugated primary antibodies prepared in blocking solution (3% goat serum and 0.3 % Triton X-100 in PBS). Primary antibodies were used as follows: Alexa Fluor 594-F4/80 (Biolegend; 123140, 1:100), Rat anti-mouse APC-CD86 (BD Pharmigen; 558703, 1:50), Alexa Fluor 647-CD206 (Biolegend; 141712, 1:100) and Alexa Fluor 488-conjugated Isolectin GS-IB_4_ (Invitrogen; I21411, 1:100). Images were acquired at higher magnification (40X and 60X) with Leica Stellaris confocal microscope. Pearson’s correlation coefficient for co-localization of F4/80 with CD86 or CD206 was calculated using Just Another Colocalization plugin (JACoP) in Image J software.

### Immunofluorescent staining of Human tissue neurofibromas

Paraffin-waxed human neurofibroma sections were stained as previously described with slight modifications(37). Briefly, to retrieve antigen epitopes, heat-mediated antigen retrieval (target retrieval solution, Dako) was performed. Sections were then rinsed in PBS-T (1X PBS with 0.25% Triton X-100), blocked in goat/donkey serum for 1 hour. Slides were incubated with primary antibodies; mouse anti-mouse CD68 (Santa Cruz; sc-17832, 1:100), and rabbit anti-mouse P-Akt2 (S474) (CST; 8599S, 1:100) overnight at 4°C on shaker. The following day, slides were washed and incubated with Alexa Fluor-conjugated secondary antibodies; goat anti-mouse Alexa Fluor 568 (Invitrogen A-21144, 1:300) and goat anti-rabbit Alexa Fluor 647 (Invitrogen; A-21244, 1:300) for 1 hour at room temperature, respectively. In between primary antibody incubations, slides were blocked again with blocking buffer for 1 hour to inhibit non-specific binding. Slides were then incubated with primary antibody; rabbit anti-mouse GLUT-1 (CST; 12939, 1:100) overnight and the next day, fluorescent secondary antibody corresponding to GLUT-1 (Alexa Fluor 488 donkey anti-rabbit; Invitrogen; A21206, 1:300) was added and incubated for 1 hour at room temperature. Nuclei were visualized by mounting medium with DAPI (Vector Laboratories; H-1200). Images were acquired at higher magnification (40X) with Leica Stellaris confocal microscope. A representative image of at least 4 patients were shown.

### Study approvals

#### Animals

All animal studies were performed in compliance with the Association for Research in Vision and Ophthalmology (ARVO) Statement for the Use of Animals in Ophthalmic and Vision Research and were approved by the Institutional Animal Care and Use Committee of Augusta University (Augusta, GA).

#### Human patient samples

The NF1 patient tissues were diagnosed and collected at Emory University. All surgeries were performed following the relevant guidelines and regulations at Emory University. The approval number of the present study by ethics committee is STUDY00002544. Written consent was obtained from all patients. After each surgery, the tissues were harvested, post-fixed in 4% paraformaldehyde overnight, and cut into blocks for paraffin embedding. The tissues were cut at 8um thickness and used for further analysis.

### Statistical Analysis

P-values were calculated by One-way ANOVA followed by Tukey’s or Sidak’s or Dunnette’s post hoc test for multiple comparisons and two-tailed Student’s t-test was used to compare pairs of data groups. In all figures, data are expressed as mean ± SD values. Figures representing *in vitro* technical replicates are presented as mean ± SEM values; (*P < 0.05, **P < 0.01, *** P < 0.001, **** P < 0.0001, ns, not significant.

## Author’s contribution

YZ and BKS designed the experiments and drafted the initial manuscript. YZ was the lead contributor to experiments, analyzed data, and created figures. RT, NZ, FN, SAHZ, RR, FZH, ZB, SK, and RB performed additional experiments and/or assisted with data analysis and interpretation. KL and NMB provided human tumor samples and assisted with data interpretation. YH, GC, EJBC, DFJ, RHK, RBC assisted with experimental design and data interpretation. All authors revised and approved the final manuscript.

## Acknowledgements

The authors acknowledge Varsha Tandra, PhD for her technical assistance.

---

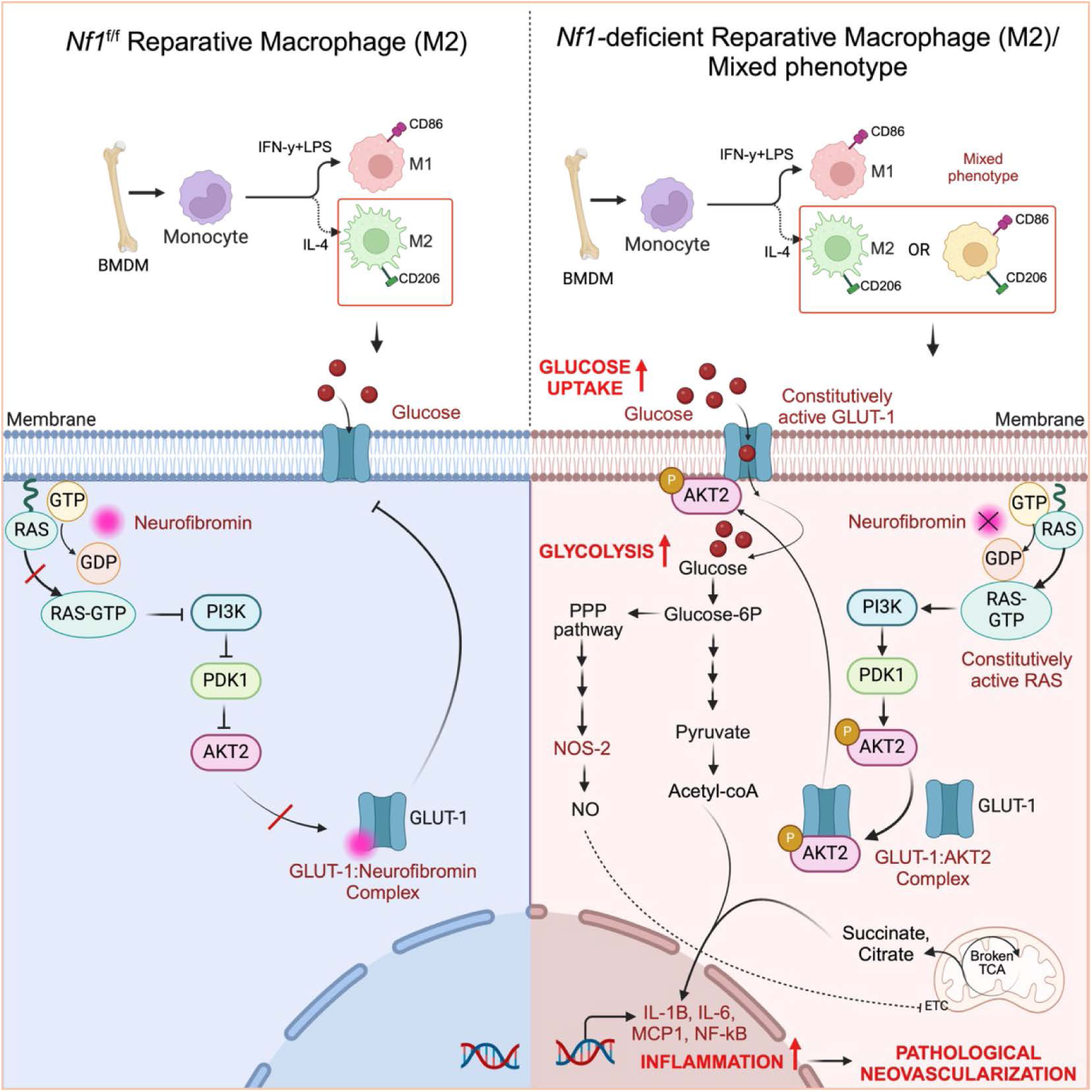

