## Supplemental Figures for "Loss of Neurofibromin Induces Inflammatory Macrophage Phenotypic Switch and Retinal Neovascularization via GLUT1 Activation"

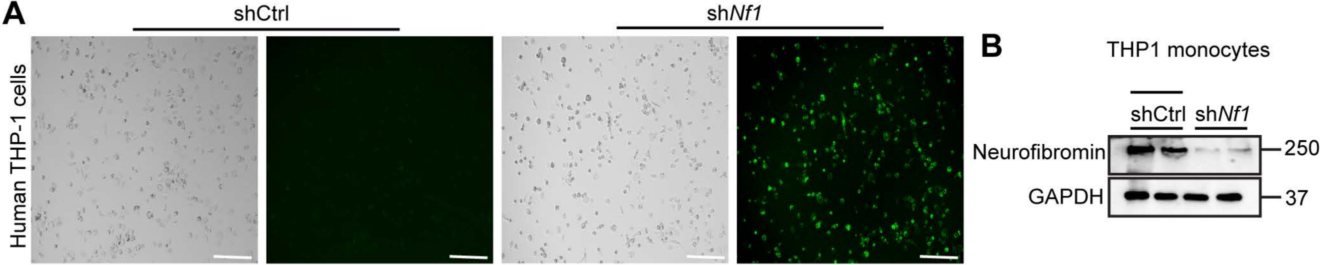


**Figure S1. Knockdown efficiency of *Nf1* in human THP1 monocytes.** (**A**) Representative fluorescent microscope images showing human THP1 monocytes transduced (GFP green signal) with control shRNA (shCtrl) and *Nf1* shRNA (sh*Nf1*) for 16 hours, followed by puromycin selection for 3 days. After puromycin selection, THP1 monocytes were differentiated by PMA (200nM) and polarized using 20ng/mL LPS and IFN-γ (for inflammatory macrophages; M1) or 20ng/mL IL-4 (for reparative macrophages; M2).Scale bars: 100μm.(**B**) Representative immunoblot showing confirmation of Nf1 knockdown in human THP1 monocytes after tranduction with shCtrl and sh*Nf1*.

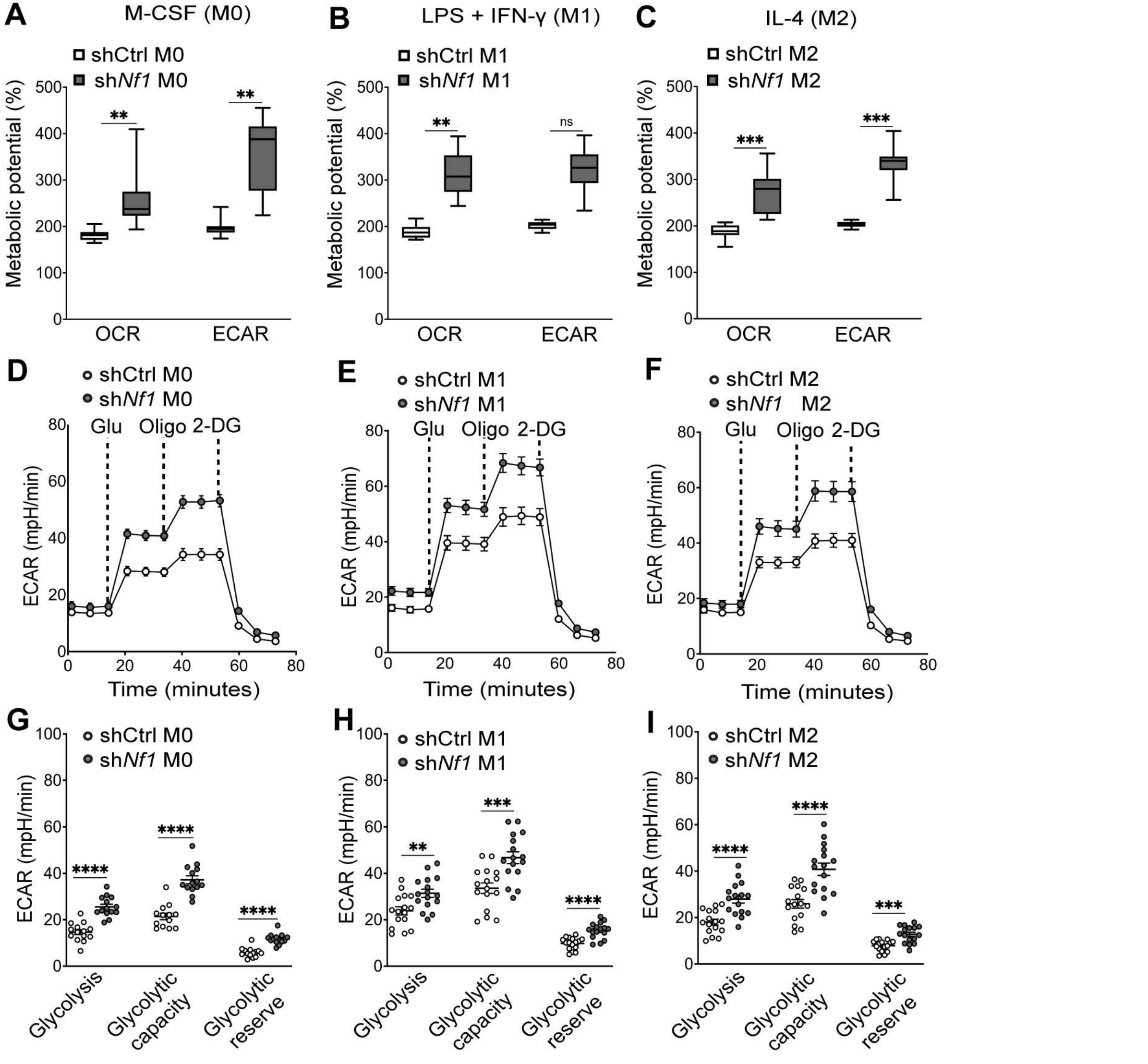


**Figure S2. *Nf1* knockdown increased glycolysis in human THP1 macrophages.** Representative measurements of (**A-C**) percent metabolic potential, and (**D-I**) Extracellular acidification rate (ECAR) performed on THP1 macrophages tranduced with shCtrl and sh*Nf1* RNA , differentiated with PMA (200nM) and polarized with 20ng/mL LPS and IFN-ϒ or 20ng/mL IL-4 for inflammatory (M1) or reparative (M2) macrophages for 16 hours, respectively. Subsequent addition of glucose, the ATP synthase inhibitor oligomycin, and the hexokinase inhibitor 2-deoxy-glucose (2-DG) were carried out where indicated (**D-F**), n = 14 technical replicates/condition. Data are expressed as mean ± SEM. P values were calculated using One-way ANOVA followed by Dunnett's T3 multiple comparision test (*P < 0.05, ** P < 0.01 , *** P < 0.001,**** P < 0.0001, ns, not significant).

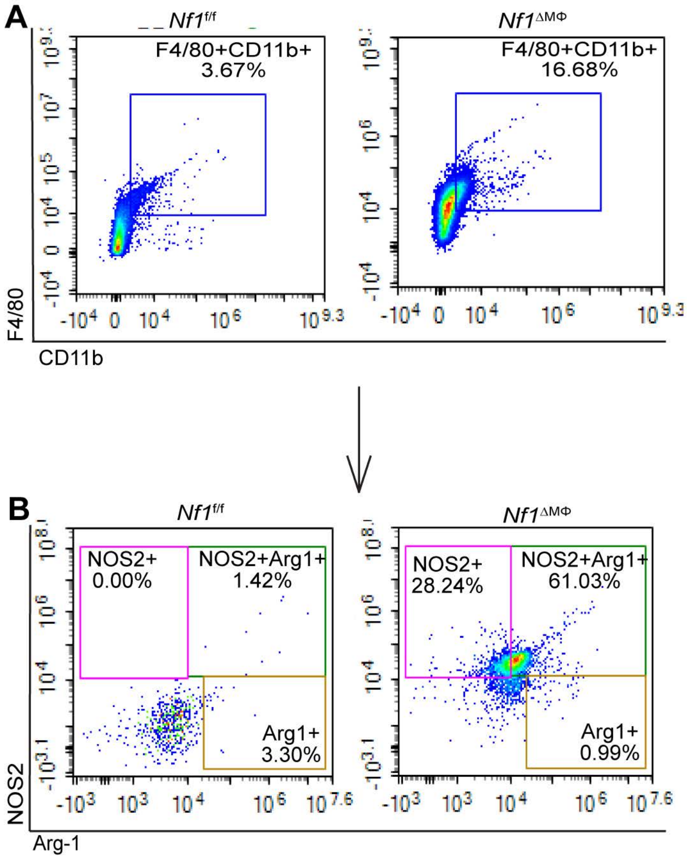


**Figure S3. Macrophage/microglia in P17 OIR retinas from *Nf1*$^{f/f}$ and *Nf1*$^{\Delta M\Phi}$ pups exhibit mixed phenotype (M1/M2-like).**
(**A**) Representative dot plots shows gating of CD11b+/F4/80+ macrophage/microglia in P17 OIR retinas.
(**B**) The selected CD11b+/F4/80+ cells were analyzed for intracellular markers NOS2 (for inflammatory or M1) and Arginase1 (reparative or M2) by flowcytometry. n= 5-6 pups/genotype, total 10-12 retinas in each genotype.
