## Supplemental Table 1 for "Loss of Neurofibromin Induces Inflammatory Macrophage Phenotypic Switch and Retinal Neovascularization via GLUT1 Activation"

**Table 1**: Molecular docking of GLUT1 with NF1 and AKT2 by H-DOCK

| **Neurofibromin** | | | **GLUT1** |  | **GLUT1** | **AKT2** | |
| --- | --- | --- | --- | --- | --- | --- | --- |
|  | GLU2123A | | VAL173 |  | THR60 | ARG371 | Protein Kinase Domain |
|  | GLU2123A | | LEU176 |  | THR63 | ARG371 |  |
|  | GLU2123A | | TYR308 |  | LEU64 | ARG371 |  |
|  | GLU2123A | | ILE311 |  | LEU67 | ARG243 |  |
|  | THR2124A | | VAL173 |  | VAL74 | ARG347 |  |
|  | THR2124A | | ILE179 |  | ILE78 | ARG245 |  |
|  | GLN2126A | | LEU169 |  | ILE78 | GLY346 |  |
|  | VAL2127A | | ILE170 |  | ILE78 | ARG347 |  |
|  | VAL2127A | | VAL173 |  | GLY79 | ARG245 |  |
|  | VAL2127A | | PHE174 |  | PHE81 | TYR351 |  |
|  | LEU2130A | | VAL166 |  | PHE81 | GLN353 |  |
|  | LEU2130A | | ILE170 |  | SER82 | ARG245 |  |
|  | GLU2134A | | LEU199 |  | LEU85 | PHE238 |  |
|  | LEU2137A | | MET13 |  | LEU85 | SER242 |  |
|  | LEU2137A | | GLY17 |  | PHE86 | SER242 |  |
|  | PRO2138A | | GLY10 |  | PHE86 | ARG243 |  |
|  | PRO2138A | | MET13 |  | PHE86 | ARG245 |  |
|  | PRO2138A | | LEU14 |  | ARG89 | PHE239 |  |
|  | LYS2139A | | LEU14 |  | PHE90 | PHE239 |  |
|  | LYS2139A | | LEU21 |  | MET98 | ARG243 |  |
|  | LYS2139A | | LEU199 |  | MET99 | SER242 |  |
|  | LYS2139A | | VAL203 |  | MET99 | ARG243 |  |
|  | PHE2140A | | LEU14 |  | MET99 | ARG245 |  |
|  | PHE2140A | | LEU199 |  | LEU101 | ARG243 |  |
|  | LEU2142A | | LEU14 |  | LEU101 | GLU244 |  |
|  | LEU2143A | | LEU199 |  | LEU102 | ARG243 |  |
|  | LEU2143A | | ILE202 |  | LEU102 | GLU244 |  |
|  | THR2179A | | ILE202 |  | LEU102 | ARG245 |  |
|  | SER2180A | | ILE202 |  | GLU120 | ARG371 |  |
|  | GLU2182A | | LEU198 |  | GLU120 | THR372 |  |
|  | GLU2182A | | ILE202 |  | ILE123 | THR372 |  |
|  | THR2183A | | LEU198 |  | LEU124 | VAL246 |  |
|  | THR2183A | | LEU199 |  | LEU124 | THR372 |  |
|  | THR2183A | | ILE202 |  | PHE127 | ARG245 |  |
|  | GLU2186A | | PHE194 |  | PHE127 | CYS345 |  |
|  | GLU2186A | | ILE195 |  | PHE127 | GLY346 |  |
|  | GLU2186A | | LEU198 |  | PHE127 | ARG347 |  |
|  | GLU2190A | | PHE174 |  | ILE128 | GLU244 |  |
|  | GLU2190A | | ILE195 |  | ILE128 | ARG245 |  |
|  | ARG2197A | | SER178 |  | ILE128 | VAL246 |  |
|  | ARG2197A | | ILE179 |  | VAL131 | ARG245 |  |
|  | ARG2197A | | MET180 |  | TYR132 | ARG245 |  |
|  | ARG2197A | | LEU185 |  | LEU135 | ARG245 |  |
|  | LYS2279A | | LEU198 |  | ILE259 | GLN353 |  |
|  | PRO2282A | | PHE112 |  | LEU260 | GLN353 |  |
|  | LEU2283A | | PHE112 |  | LEU260 | HIS355 |  |
|  | GLU2318A | | LEU101 |  | PHE263 | GLN353 |  |
|  | HIS2322A | | LEU101 |  | PHE263 | ASP354 |  |
|  | HIS2322A | | PHE104 |  | ARG264 | ASP354 |  |
|  | HIS2322A | | VAL105 |  | ARG264 | HIS355 |  |
|  | THR2323A | | VAL108 |  | ARG264 | GLU356 |  |
|  | SER2326A | | VAL108 |  | ALA405 | GLN353 |  |
|  | SER2326A | | LEU109 |  | VAL406 | GLN353 |  |
|  | SER2326A | | MET121 |  | PHE409 | TYR351 |  |
|  | SER2381A | | PRO205 |  | PHE409 | ASN352 |  |
|  | PRO2382A | | ARG93 |  | PHE409 | GLN353 |  |
|  | PRO2382A | | ASN94 |  | ARG89 | TYR438 | AGC-Kinase C Terminal Domain |
|  | PRO2382A | | GLU209 |  | ARG89 | PHE439 |  |
|  | ALA2383A | | ARG93 |  | ARG89 | ASP440 |  |
|  | ALA2383A | | ASN94 |  | ARG89 | ASP441 |  |
|  | ALA2383A | | LEU97 |  | PHE90 | ARG437 |  |
|  | ALA2383A | | PRO205 |  | PHE90 | ASP441 |  |
|  | ALA2386A | | ASN94 |  |  | | |
|  | ALA2386A | | MET98 |  |  |  |  |
|  | VAL2389A | | MET98 |  |  |  |  |
|  | ARG2390A | | LEU97 |  |  |  |  |
|  | ARG2390A | | MET98 |  |  |  |  |
|  | ARG2390A | | MET99 |  |  |  |  |
|  | ARG2390A | | LEU101 |  |  |  |  |
|  | VAL2424A | | ARG89 |  |  |  |  |
|  | VAL2424A | | PHE90 |  |  |  |  |
|  | MET739B | | LYS255 |  |  |  |  |
|  | GLU740B | | ARG253 |  |  |  |  |
|  | SER743B | | ARG253 |  |  |  |  |
|  | VAL744B | | ARG253 |  |  |  |  |
|  | LYS757B | | ARG249 |  |  |  |  |
|  | ARG758B | | ARG249 |  |  |  |  |
|  | ARG758B | | ARG253 |  |  |  |  |
|  | ALA761B | | ARG249 |  |  |  |  |
|  | ALA761B | | ARG253 |  |  |  |  |
|  | LEU762B | | ARG253 |  |  |  |  |
|  | ARG765B | | GLN250 |  |  |  |  |
|  | ARG765B | | ARG253 |  |  |  |  |
| Ras GAP  Domain  (CHAIN B) | ARG1375B | | ILE339 |  |  |  |  |
|  | ARG1375B | | ALA342 |  |  |  |  |
|  | ARG1375B | | PHE378 |  |  |  |  |
|  | PHE1376B | | PHE378 |  |  |  |  |
|  | PRO1377B | | PHE320 |  |  |  |  |
|  | GLN1378B | | VAL316 |  |  |  |  |
|  | GLN1378B | | PHE320 |  |  |  |  |
|  | GLN1378B | | VAL370 |  |  |  |  |
|  | GLN1378B | | PHE373 |  |  |  |  |
|  | GLN1378B | | GLY374 |  |  |  |  |
|  | GLN1378B | | ALA377A |  |  |  |  |
|  | ASN1379B | | VAL370A |  |  |  |  |
|  | HIS1431B | | TYR366A |  |  |  |  |
|  | VAL1432B | | TRP363A |  |  |  |  |
|  | LEU1433B | | TRP363A |  |  |  |  |
|  | PHE1434B | | TRP363A |  |  |  |  |
|  | THR1435B | | LEU361A |  |  |  |  |
|  | THR1435B | | TRP363A |  |  |  |  |
|  | ARG1441B | | GLN360A |  |  |  |  |
|  | ARG1441B | | LEU361A |  |  |  |  |
|  | ARG1441B | | PRO362A |  |  |  |  |
