## Supplemental Table 2 for "Loss of Neurofibromin Induces Inflammatory Macrophage Phenotypic Switch and Retinal Neovascularization via GLUT1 Activation"

**Supplemental table 2.** Summary of human NF1 patients selected for neurofibromas tissues sectioning for immunofluorescent staining of macrophages, GLUT1 and P-Akt2.

| **Study ID** | **Age (Years)** | **Sex** | **NF1** | **Location** |
| --- | --- | --- | --- | --- |
| NF1-12 | 27 | F | Y | L Brachial Plexus |
| NF1-19 | 32 | M | Y | R Posterior Thigh |
| NF1-21 | 38 | F | Y | R Mid-Calf |
| NF1-26 | 38 | F | Y | C3-C4 Intradural |
| NF1-23 | 27 | M | Y | R Retroperitoneal |
