## Supplemental Table 3 for "Loss of Neurofibromin Induces Inflammatory Macrophage Phenotypic Switch and Retinal Neovascularization via GLUT1 Activation"

**Supplemental table 3: Primers for qRT-PCR used in the study:**

| Mice Gene ID | Sequence / Reference ID | Probes | Source |
| --- | --- | --- | --- |
| *MCP-1* | Forward: GGCTCAGCCAGATGCAGTTAA  Reverse: CCTACTCATTGGGATCATCTTGCT | SYBR Green | IDT |
| *NOS2* | Forward: GGCAGCCTGTGAGACCTTTG  Reverse: TGCATTGGMGTGAAGCGTTT | SYBR Green | IDT |
| *IL-10* | Forward: GCTCTTACTGACTGGCATGAG  Reverse: CGCAGCTCTAGGAGCATGTG | SYBR Green | IDT |
| *IL-1β* | Forward: TGCCACCTTTTGACAGTGATG  Reverse: ATGTGCTGCTGCGAGATTTG | SYBR Green | IDT |
| *NFkB (Rela (p65)* | Forward: TCCTGTTCGAGTCTCCATGCAG  Reverse: GGTCTCATAGGTCCTTTTGCGC | SYBR Green | IDT |
| *Tnfa* | Forward: GGTCCCCAAAGGGATGAGAA  Reverse: TGAGGGTCTGGGCCATAGAA | SYBR Green | IDT |
| *Rplp0* | Forward: CCTCCTTCTTCCAGGCTTTG  Reverse: CCACCTTGTCTCCAGTCTTTATC | SYBR Green | IDT |
| *Arginase-1* | Mm00475988-m1 | FAM | TaqMan Assay |
| *IL-6* | Mm00446190-m1 | FAM | TaqMan Assay |
| *GLUT1* | Forward: GCAGTTCGGCTATAACACTGG  Reverse: GCGGTGGTTCCATGTTTGATT) | SYBR Green | IDT |
